# Kinesin-Binding Protein (KBP) buffers the activity of KIF18A and KIF15 in mitosis to ensure accurate chromosome segregation

**DOI:** 10.1101/343046

**Authors:** Heidi L. H. Malaby, Megan E. Dumas, Ryoma Ohi, Jason Stumpff

## Abstract

Mitotic kinesins must be regulated to ensure a precise balance of spindle forces and accurate segregation of chromosomes into daughter cells. Here we demonstrate that Kinesin-Binding Protein (KBP) reduces the activity of KIF18A and KIF15 during metaphase. Overexpression of KBP disrupts the movement and alignment of mitotic chromosomes and decreases spindle length, a combination of phenotypes observed in cells deficient for KIF18A and KIF15, respectively. We show through gliding filament and microtubule co-pelleting assays that KBP directly inhibits KIF18A and KIF15 motor activity by preventing microtubule-binding. Consistent with these effects, the mitotic localizations of KIF18A and KIF15 are altered by overexpression of KBP. Cells depleted of KBP exhibit lagging chromosomes in anaphase, an effect that is recapitulated by KIF15 and KIF18A overexpression. Based on these data, we propose a model in which KBP acts as a protein buffer in mitosis, protecting cells from excessive KIF18A and KIF15 activity to promote accurate chromosome segregation.

**SUMMARY:** Kinesin-Binding Protein (KBP) is identified as a regulator of the kinesins KIF18A and KIF15 during mitosis. KBP buffers the activity of these motors to control chromosome alignment and spindle integrity in metaphase and prevent lagging chromosomes in anaphase.

## INTRODUCTION

Stochastic variations in gene transcription within individual isogenic cells lead to non-uniform protein levels on a cell-to-cell basis (Sigal et al., 2006). These in turn can affect the rate and efficiency of all physiological processes, necessitating counter-measures to buffer the cell against alterations in protein levels that would otherwise be detrimental. Mitosis is particularly sensitive to biological variations in protein expression levels: abnormally high or low concentrations of mitotic regulators can lead to errors in mitotic spindle function and chromosome segregation. Given the importance of force-balance within the mitotic spindle for its assembly and function, it is clear that mechanisms to regulate the activities of molecular motors, such as the mitotic kinesins, would be important for cell division. Indeed, too much or too little mitotic kinesin activity can impair mitotic progression. For example, loss of KIF18A (kinesin-8) function leads to chromosome alignment defects and abnormally long mitotic spindles, whereas cells with increased KIF18A levels form short or multipolar spindles (Du et al., 2010; Mayr et al., 2007; Stumpff et al., 2008). Similarly, increasing or decreasing MCAK (kinesin-13) leads to abnormal chromosome movements and kinetochore-microtubule attachments (Wordeman et al., 2007). Thus, mitosis requires regulatory mechanisms that promote optimal levels of motor activity within the spindle.

Sequestration and inactivation of kinesins is one possible mechanism to acutely and reversibly regulate motor activity levels, and kinesin-binding protein (KBP) appears to fulfill this role in at least some cellular contexts. KBP was first identified as a disease-causing gene (dubbed *KIAA1279*) through genetic sequencing of patients with Goldberg-Shprintzen Syndrome, a disorder marked by microcephaly, mental retardation, and facial dysmorphism (Alves et al., 2010; Brooks et al., 2005; Drevillon et al., 2013). In neurons, KBP directly inhibits the kinesin-3 family member Kif1Bα, which is necessary for proper mitochondrial transport down the axon (Wozniak et al., 2005). Mitochondrial derived acetyl-CoA can activate signaling leading to the acetylation of KBP, which is then targeted for degradation, a process linked to decreased mitochondrial biogenesis (Donato et al., 2017). Recently, an extensive biochemical study reported KBP-specific binding to a large panel of human kinesins (Kevenaar et al., 2016). In addition to kinesin-3 family members (KIF1A, KIF1B, KIF1C, and KIF14), KBP was also found to have strong affinity for kinesin-2 family members (KIF3A and KIF3B), a kinesin-8 (KIF18A), and a kinesin-12 (KIF15). KBP interacts directly with kinesin motor domains with a 1:1 KBP:motor domain stoichiometry. Motor domain-binding by KBP inhibits the kinesin-microtubule interaction (Kevenaar et al., 2016).

The discovery that KBP binds KIF18A and KIF15 suggests that KBP may also play an important role in regulating cell division, as both of these motors have established mitotic functions. KIF18A localizes to the ends of kinetochore-microtubules and confines chromosomes to the spindle equator by regulating kinetochore microtubule dynamics (Du et al., 2010; Stumpff et al., 2008). KIF15 maintains spindle integrity by modulating kinetochore-microtubule separation forces (Sturgill and Ohi, 2013; Tanenbaum et al., 2009). Consistent with a role for KBP in regulating the mitotic functions of KIF15 and KIF18A, it was identified as an interactor of both motors in cycling HeLa cells (Maliga et al., 2013) and has been implicated in regulating KIF15’s functions within the spindle (Brouwers et al., 2017).

Here, we establish that KBP inhibits the functions of both KIF18A and KIF15 to promote proper spindle function during mitosis. KBP directly binds the motor domains of KIF18A and KIF15, inhibiting their interactions with microtubules. Accordingly, increased expression of KBP, which is primarily cytosolic, disrupts the normal localization of both KIF18A and KIF15 within the spindle. KBP is required to ensure proper chromosome segregation during anaphase, and we show that an excess of either KIF18A or KIF15 results in similar chromosome segregation defects. Based on these data, we propose that KBP functions to buffer kinesin activity during mitosis, ensuring that the activities of KIF15 and KIF18A are maintained within a zone compatible with accurate chromosome segregation.

## RESULTS

### KBP expression level affects chromosome alignment and spindle length in mitotic cells

To investigate a potential role for KBP during mitosis, we examined KBP’s localization and the effects of altering its expression in dividing cells. HeLa and RPE1 cells were transfected with either a control siRNA or a previously validated KBP-specific siRNA (Brouwers et al., 2017; Kevenaar et al., 2016). Immunoblotting of cell extracts showed that a specific band near 75 kDa was detected with anti-KBP antibodies and that this band was reduced following KBP siRNA treatment (Figure 1A and Figure S1). This is consistent with the predicted molecular weight of KBP (72 kDa) and previous findings (Drevillon et al., 2013; Kevenaar et al., 2016; Wozniak et al., 2005). Endogenous KBP has been reported to localize to the mitotic spindle in metaphase cells by immunofluorescence (Brouwers et al., 2017). While we also detected spindle staining with an anti-KBP antibody, it was largely unchanged after KBP siRNA treatment in HeLa and RPE1 cells (Figure S1), suggesting that spindle staining with the KBP antibody is non-specific. Conversely, an N-terminally mCherry-tagged KBP construct was primarily cytosolic with only faint centrosome and spindle staining (Figure 1B, 1C, and S1). In interphase cells expressing high levels of mCherry-KBP, KBP was also cytosolic with occasional puncta co-localizing with microtubules (Figure 1B, inset). These findings support previous studies that KBP itself does not exhibit robust microtubule-binding activity (Kevenaar et al., 2016).

**Figure 1.**
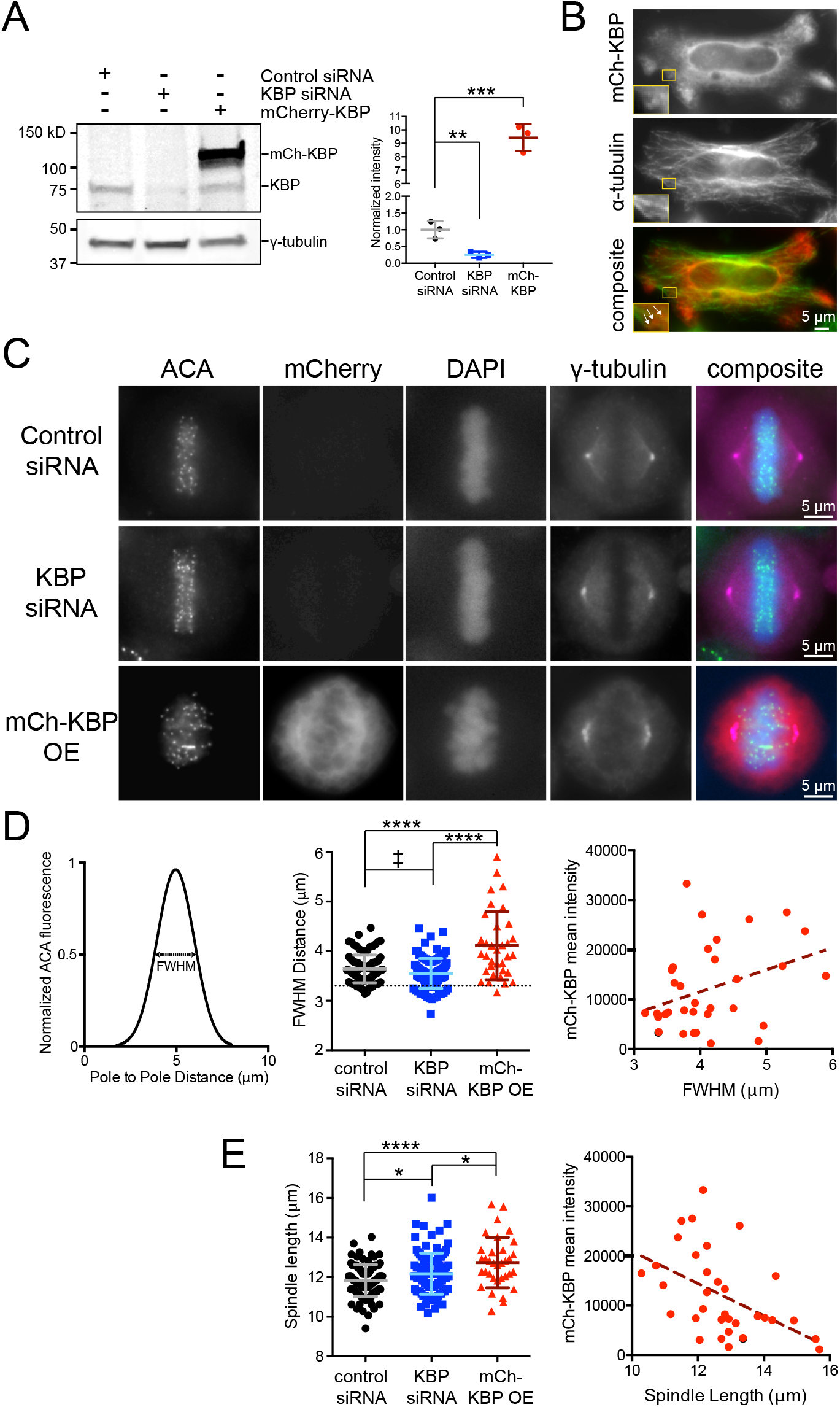
KBP regulates chromosome alignment and spindle length in mitotic cells. (**A**) Western Blot against KBP, showing protein levels under KBP siRNA and mCherry-tagged (mCh) KBP transfection conditions from HeLa cells. γ-tubulin was used as a loading control. (*right*) KBP knockdown and mCh-KBP overexpression quantification, obtained from quantification of three independent blots, corrected against γ-tubulin and normalized to the control siRNA transfection. **, p = 0.009, ***, p = 0.0002 by unpaired t-test comparing each condition to control siRNA. (**B**) mCh-KBP does not bind microtubules in interphase HeLa cells. Yellow box denotes inset area. Arrows highlight occasional mCh-KBP puncta that co-localize with α-tubulin. (**C**) Representative metaphase HeLa cells arrested in MG132 were treated with control or KBP siRNAs or overexpress (OE) mCh-KBP. ACA, anti-centromere antibodies. (**D**) Chromosome alignment was quantified by determining the full-width at half maximum (FWHM) of a Gaussian fit to the distribution of ACA fluorescence along the spindle axis. (*left*) Graphical representation of FWHM measurement. (*middle*) FWHM distance values for each cell under the indicated conditions. Dotted line denotes cutoff value for hyper-aligned cells (3.3 μm), empirically determined from the control population. ‡, p = 0.0432 by Chi-square analysis comparing hyper-aligned populations. ****, adjusted p < 0.0001 with 95% confidence interval by one-way ANOVA analysis with Tukey’s multiple comparisons test of full data sets. (*right*) Correlation plot of mCh-KBP fluorescence intensity vs. FWHM alignment values. Dotted line is linear regression to show data trend. (**E**) (*left*) Plot of spindle lengths measured in cells following the indicated treatments. *, adjusted p < 0.05, ****, adjusted p < 0.0001 with 95% confidence interval by one-way ANOVA with Tukey’s multiple comparisons test. (*right*) Correlation plot of mCh-KBP fluorescence intensity vs. spindle lengths. Dotted line is linear regression to show data trend. Data in (D) and (E) obtained from three independent experiments with the following cell numbers: control siRNA (96), KBP siRNA (105), mCh-KBP OE (34).

To examine the effects of KBP on early mitotic events, HeLa and RPE1 cells were transfected with either KBP siRNAs or mCherry-KBP, arrested in MG132 to prevent entry into anaphase, fixed, and stained to visualize chromosomes, centromeres, centrosomes, and microtubules (Figure 1C). Increasing or decreasing KBP levels led to aberrations in chromosome alignment and spindle length in metaphase cells. Chromosome alignment was quantified by measuring centromere distribution along the spindle axis and using the full width at half of maximum (FWHM) as a metric for comparison across cell populations and treatment conditions (Kim et al., 2014; Stumpff et al., 2012). KBP siRNA treatment significantly increased the proportion of metaphase cells displaying hyper-alignment, characterized by a bilinear arrangement of sister kinetochores at the metaphase plate (Figure 1D, middle and Figure S2A-B). Conversely, mCherry-KBP overexpression significantly decreased chromosome alignment. This effect was dose-dependent, as chromosome alignment scaled proportionally with mCherry intensity in both HeLa and RPE1 cells (Figure 1D, right and Figure S2C). Alterations in KBP expression also affected spindle lengths. KBP siRNA treatment resulted in a population of slightly longer spindles (Figure 1E and Figure S2D). Interestingly, mCherry-KBP overexpression also increased average spindle length, but higher KBP expression led to shorter spindles in both HeLa and RPE1 cells (Figure 1E and S2D). Similar results were obtained in asynchronously dividing HeLa cells that were not treated with MG132 (Figure S3). Together, these results support a role for KBP in regulating chromosome alignment and spindle length during mitosis.

### Co-depletion of KIF18A and KIF15 recapitulates the spindle length and chromosome alignment defects seen in KBP overexpressing cells

KBP was previously identified as a potential interactor of KIF18A, KIF15, and KIF14 from affinity-purification/mass spectrometry analysis of mitotic kinesins (Maliga et al., 2013), and these three mitotic motors were also characterized as KBP substrates by co-IPs (Kevenaar et al., 2016). While KIF18A’s influence of chromosome alignment (Du et al., 2010; Stumpff et al., 2008) and KIF15’s effects on spindle length (Sturgill and Ohi, 2013; Tanenbaum et al., 2009) are well established, a role for KIF14 in metaphase has not been documented. KIF14 was cytosolic in metaphase cells and localized to the midbody during telophase (Figure S4A). KIF14-specific siRNA treatment significantly reduced KIF14 immunofluorescence but did not detectably affect chromosome alignment or spindle length (Figures S4A-C). Therefore, we focused on investigating KIF18A and KIF15 as metaphase KBP substrates.

To determine the effects of KIF18A and KIF15 co-inhibition, we knocked down KIF18A, KIF15, or KIF18A and KIF15 using previously validated siRNAs (Kim et al., 2014; Stumpff et al., 2008; Stumpff et al., 2012; Sturgill et al., 2016; Sturgill and Ohi, 2013; Tanenbaum et al., 2009). Metaphase cells were then fixed and stained to visualize centromeres, centrosomes, chromosomes, and kinesins (Figure 2A and 2B). Chromosome alignment and spindle length were analyzed as in Figure 1. KIF18A knockdown resulted in unaligned chromosomes (large FWHM values), and this phenotype remained unchanged in combination with KIF15 knockdown (Figure 2C, left). This was consistent with our observation that chromosomes align during metaphase in *KIF15Δ* knock-out cells (Sturgill et al., 2016 and Figure S4D-E). Consistent with previous studies, single KIF18A and KIF15 knockdowns affected spindle length, with the former increasing and the latter decreasing the average distance between spindle poles (Mayr et al., 2007; Stumpff et al., 2008; Stumpff et al., 2012; Sturgill et al., 2016; Sturgill and Ohi, 2013and Figure S4D-E). Interestingly, HeLa cells co-depleted of KIF18A and KIF15 displayed spindle lengths on par with control cells (Figure 2C, right). KIF18A and KIF15 depletion was confirmed by immunofluorescence (Figure 2D). These results indicate that inhibition of KIF18A and KIF15 in combination could generate the effects observed upon KBP overexpression in Figure 1. We also note that slightly longer spindle lengths observed in cells at lower mCherry-KBP expression levels could be explained by an increased ability of KBP to inhibit KIF18A compared with KIF15 (Figure 1E).

**Figure 2.**
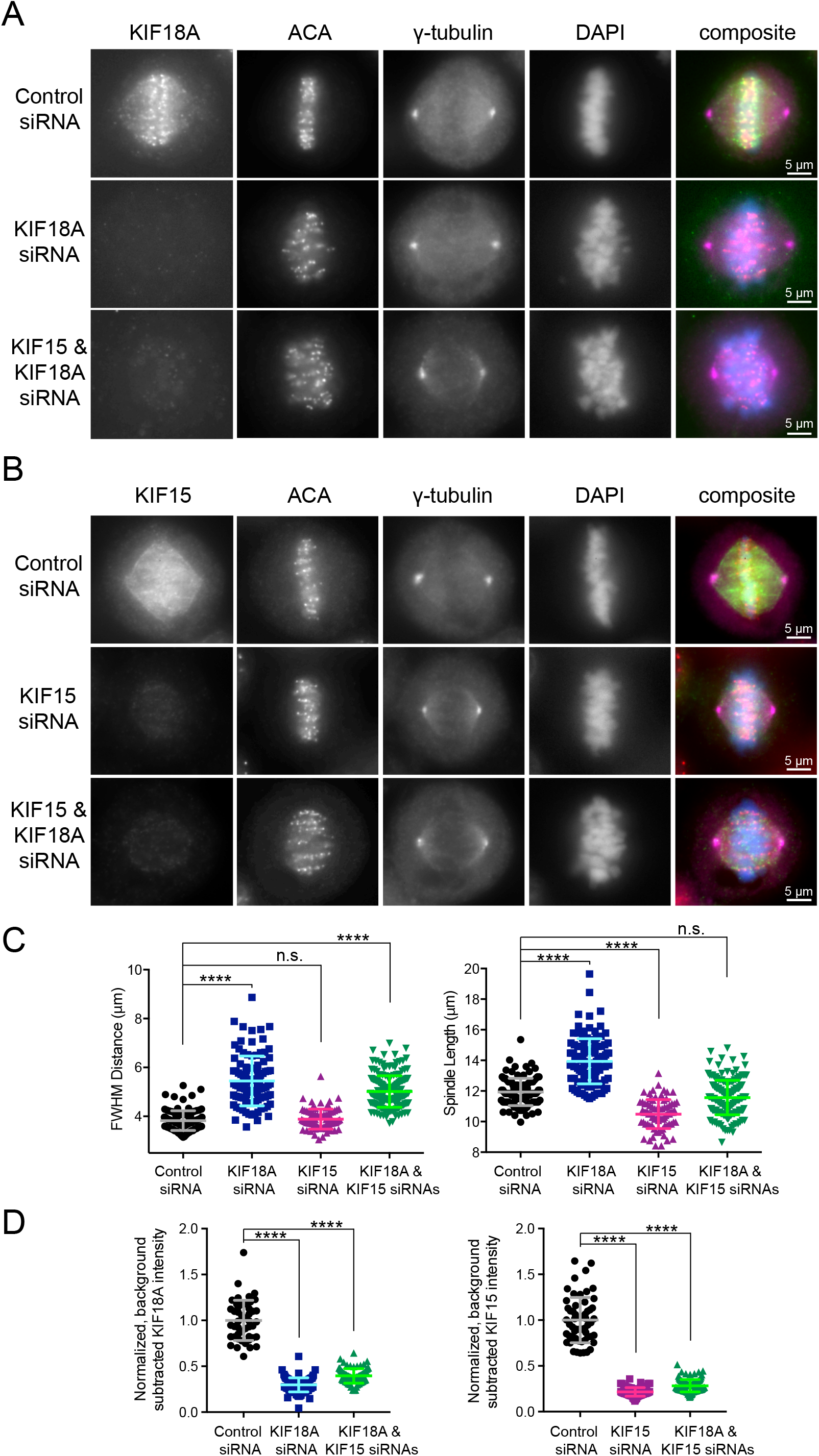
Metaphase cells deficient in KIF18A and KIF15 display chromosome alignment and spindle length abnormalities. Metaphase cells arrested in MG132 with antibody staining for (**A**) endogenous KIF18A or (**B**) endogenous KIF15. ACA, anti-centromere antibodies. (**C**) Quantification of FWHM (*left*) and spindle length (*right*) for individual cells treated with the indicated siRNAs. n.s., not significant; ****, adjusted p < 0.0001 with 95% confidence interval by one-way ANOVA with Tukey’s multiple comparisons test. (**D**) Quantification of KIF18A (*left*) and KIF15 (*right*) knockdown efficiencies for cells analyzed in (C). For the double knockdown quantification, one coverslip each was stained for each kinesin. ****, adjusted p < 0.0001 with 95% confidence interval by one-way ANOVA with Tukey’s multiple comparisons test as compared to control siRNA treatment. Data in (C) and (D) obtained from two independent experiments from the following cell numbers: control siRNA (103), KIF18A siRNA (113), KIF15 siRNA (82), KIF18A and KIF15 siRNA (184).

### KBP inhibits KIF18A motor activity more potently than KIF15 motor activity *in vitro*

To directly measure and compare the effects of KBP on KIF18A and KIF15 activity, C-terminally truncated KIF18A (KIF18A-N480-GFP-His_6_) and KIF15 (His_6_-KIF15-N700) were purified from *Sf*9 cells or *E. coli* and analyzed *via* gliding filament assays in the presence and absence of purified GST-KBP (Figure 3A and 3B, Figure S5 A and B, Videos S1-S4). Consistent with previous studies, we found that KBP inhibits the activity of both KIF15 and KIF18A (Kevenaar et al., 2016), and this inhibition occurs in a dose-dependent manner (Figure 3A and 3B), with KIF18A showing enhanced sensitivity to KBP levels (~80% inhibition of KIF18A activity achieved with 100 nM KBP compared to 250 nM KBP for KIF15).

**Figure 3.**
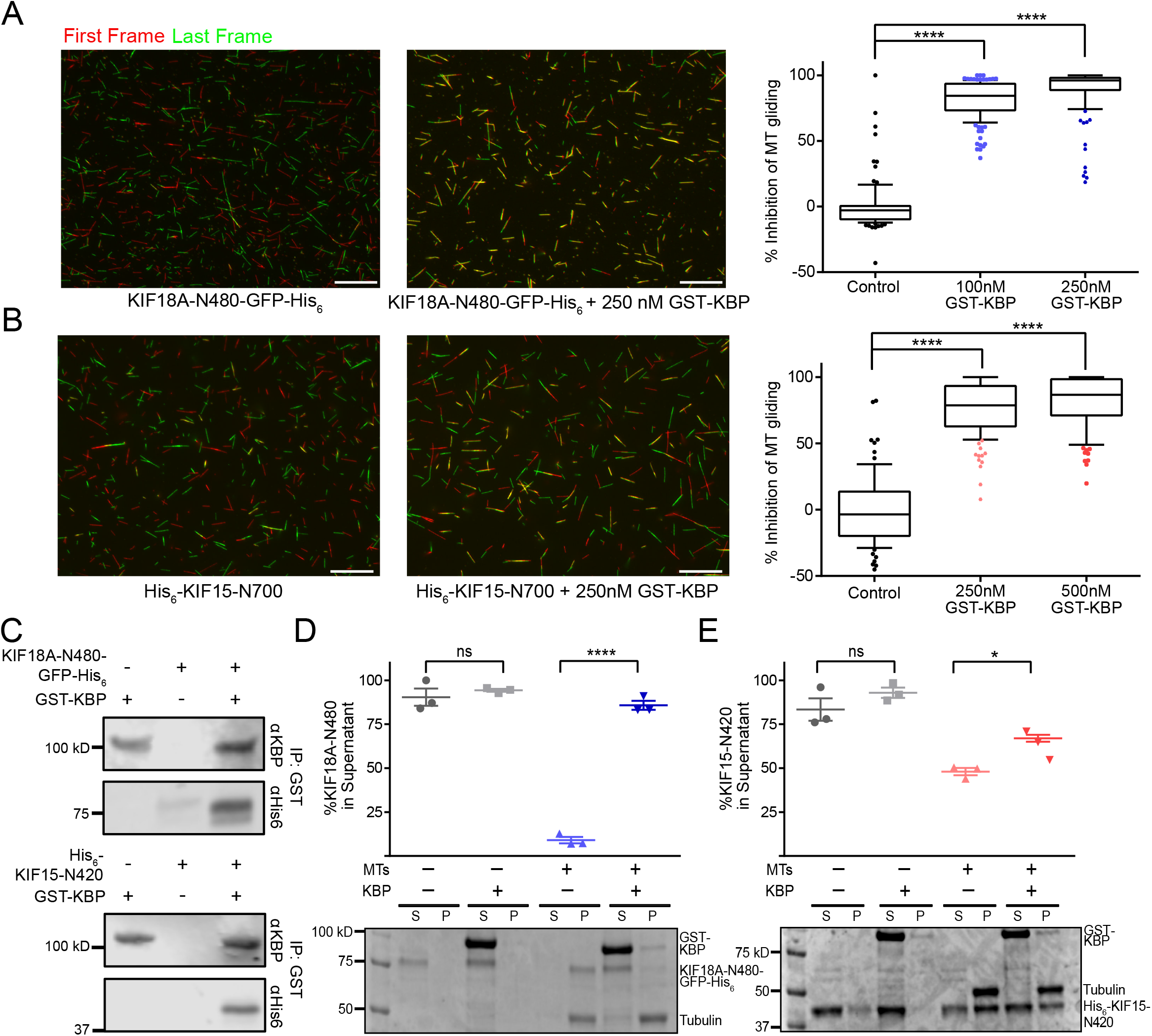
KBP inhibits KIF15 and KIF18A motor function in vitro. (**A**) Gliding filament assay with KIF18A-N480-GFP-His_6_ and GST-KBP. (*left*) Representative fields showing the first frame (red) and last frame (green) of microtubule filaments in the presence or absence of GST-KBP. Scale, 20 μm. Full movies (Videos S1 and S2) available in supplemental. (*right*) Quantification of percent inhibition of microtubule gliding for KIF18A-N480-GFP-His_6_ alone (N = 95), or with 100 nM (N = 147) or 250 nM (N = 123) GST-KBP present in the flow cell. N = total # of MTs measured from triplicate independent experiments. (**B**) Gliding filament assay with His_6_-KIF15-N700 and GST-KBP. (*left*) Representative fields showing the first frame (red) and last frame (green) of microtubule filaments in the presence or absence of GST-KBP. Full movies (Videos S3 and S4) available in supplemental. (*right*) Quantification of microtubule gliding velocity for His_6_-KIF15-N700 alone (N = 76), or with 250 nM (N = 125) or 500 nM GST-KBP (N = 92) present in the flow cell. N = total # of MTs measured from triplicate independent experiments. ****, adjusted p < 0.0001 with 95% confidence interval by one-way ANOVA with Tukey’s multiple comparisons test as compared to control gliding reactions. (**C**) Western blot showing co-immunoprecipitation (co-IP) of purified KIF18A-N480-GFP-His_6_ (50 nM) (*top*) or His_6_-KIF15-N420 (50 nM) (*bottom*) with GST-KBP (250 nM) was probed with antibodies targeting 6xHis (α His) and KBP (α KBP). (**D**) and (**E**) Results of co-pelleting assays for KIF18A-N480-GFP-His_6_ (D) and His_6_-KIF15-N420 (E) in the presence and absence of GST-KBP and/or microtubules (MTs). S, supernatant; P, pellet. Representative coomassie gels are shown along with quantification of percentage of kinesin motor present in co-pelleting fractions. Data obtained were from three independent experiments. ****, adjusted p < 0.0001, *, adjusted p = 0.031 with 95% confidence interval by one-way ANOVA with Tukey’s multiple comparisons test.

KBP inhibits microtubule binding of transport motors such as KIF1A (Kevenaar et al., 2016). To determine if this is also the case for KIF18A and KIF15, we pre-incubated motors with KBP to allow complex formation, and then measured their ability to bind microtubules in the presence of AMPPNP. The motor constructs used in these assays (KIF18A-N480-GFP-His_6_ and His_6_-KIF15-N420) lacked secondary microtubule-binding sites outside of the motor domain (Reinemann et al., 2017; Stumpff et al., 2011), ensuring that any effects of KBP on motor-microtubule interactions were likely to be mediated through the motor domain. We confirmed that KBP directly associates with both KIF18A-N480 and KIF15-N420 (Figure 3C). In the absence of KBP, both KIF18A-N480 and KIF15-N420 co-pelleted with microtubules, although with varying efficiencies (Figure 3D and 3E). In the presence of microtubules and KBP, both motors shifted from the pellet to the supernatant. (> 90% for KIF18A, ~20% for KIF15) (Figure 3D and 3E). Interestingly, although KBP also directly binds full-length KIF18A, it does not dramatically reduce the full-length motor’s ability to pellet with microtubules (Figure S5C and S5E). Consistent with our previous findings showing that KIF15 auto-inhibits its ability to bind microtubules, full-length KIF15 did not interact with microtubules *in vitro* (Sturgill et al., 2014) and also did not bind purified KBP (Figure S5D and S5F). Taken together, these data indicate that KBP directly inhibits KIF18A and KIF15 by blocking the microtubule binding activity of their motor domains but may not prevent microtubule interaction when secondary non-motor microtubule-binding elements are present.

### KBP expression alters the spindle localization of KIF18A and KIF15

To determine if KBP alters the interaction of KIF18A and KIF15 with microtubules in mitotic cells, we analyzed the localizations of endogenous KIF18A (Figure 4A-B) and KIF15 (Figure 4C-D) following KBP knockdown or overexpression. KIF18A normally accumulates at kinetochore microtubule plus-ends during metaphase. Measurements of KIF18A distribution along individual kinetochore microtubule fibers indicated that KIF18A plus-end accumulation was insensitive to KBP knockdown but that the motor showed more uniform spindle distribution following KBP overexpression (Figure 4B).

**Figure 4.**
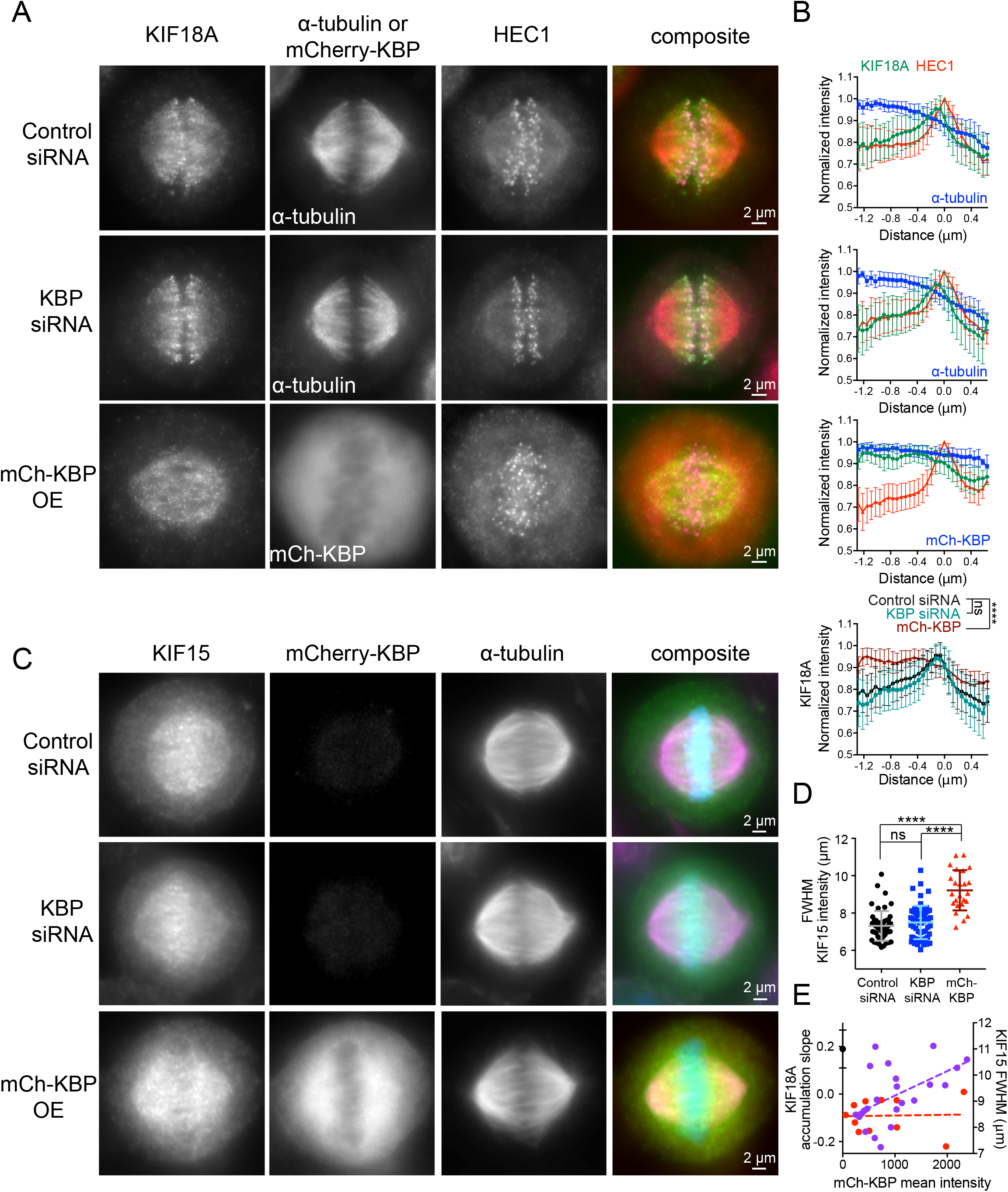
KBP overexpression alters the localization of both KIF18A and KIF15 on the mitotic spindle. (**A**) Localization of endogenous KIF18A in metaphase cells arrested in MG132 after the indicated treatments (OE = overexpression). HEC1 was used as a kinetochore marker. (**B**) Line scans measuring KIF18A distribution along kinetochore microtubules. Multiple line scans were averaged after normalizing to peak HEC1 fluorescence intensity. KIF18A, green; HEC1, red; α-tubulin, blue. Solid line indicates the mean, error bars are standard deviation. (*bottom*) Overlay of average KIF18A localization for all three conditions: control siRNA, black; KBP siRNA, blue-green; mCh-KBP, red. n.s, not significant; ****, p < 0.0001 by F-test comparing slopes of linear regressions. (**C**) Localization of endogenous KIF15 in metaphase cells under the indicated treatment conditions. (**D**) Quantification of KIF15 localization on the spindle in cells overexpressing KBP. The FWHM of KIF15 intensity was determined for each cell analyzed. n.s., not significant; ****, adjusted p < 0.0001 with 95% confidence interval by one-way ANOVA with Tukey’s multiple comparisons test. (**E**) Correlation plot of KIF15 FWHM alignment values (right y-axis, purple) and slopes of KIF18A accumulation (left y-axis, red) as a function of mCh-KBP fluorescence in individual cells. KIF18A distribution slopes were averaged from 2-4 line scans per cell. Black dot denotes average KIF18A accumulation slope under control siRNA condition with standard deviation. Dotted lines are linear regression to show data trend. Data in (B) and (D) obtained from three independent experiments each with the following cell and line scan numbers: (B) control siRNA (21 (61 lines)), KBP siRNA (22 (63 lines)), mCh-KBP OE (19 (53 lines)); (D) control siRNA (51), KBP siRNA (59), mCh-KBP OE (24).

Endogenous KIF15 localizes to microtubules throughout the spindle, with a slightly higher concentration observed near the metaphase plate (Figure 4C). To quantify changes in KIF15 localization, we determined the width of KIF15’s fluorescence distribution along the spindle axis at half of its maximum intensity (FWHM), similar to the metric used to quantify chromosome alignment described in Figure 1. KIF15 localization was not significantly altered upon KBP knockdown but was distributed more evenly along the spindle in cells overexpressing mCherry-KBP (Figure 4D). While endogenous KIF18A accumulation is abolished in even the faintest mCherry-KBP expressing cells, KIF15 distribution correlates with mCherry-KBP expression level in the cell (Figure 4E). Overall, these findings confirm that KBP affects the localization of both KIF18A and KIF15 in mitosis and further support the notion that KBP is a more potent inhibitor of KIF18A than KIF15.

### The C-termini of KIF18A and KIF15 facilitate spindle localization in the presence of mCherry-KBP

The differential effects of KBP on the microtubule co-pelleting of full-length KIF18A compared to KIF18A-N480 suggest that the microtubule-binding site located in KIF18A’s C-terminal tail may facilitate microtubule association in the presence of KBP (Mayr et al., 2011; Stumpff et al., 2011; Weaver et al., 2011). To test whether KIF18A’s spindle localization in cells overexpressing KBP depends on its C-terminus, we compared the localizations of KIF18A full-length (FL) and KIF18A-N480 tagged with GFP (Figure 5). GFP-Kif18A-FL displayed a more uniform spindle localization and an increase in cytoplasmic motor in the presence of mCherry-KBP (Figure 5B). Strikingly, mCherry-KBP expression abolished the spindle localization of GFP-KIF18A-N480, resulting in nearly complete redistribution of the motor to the cytosol (Figure 5A and 5B). This finding 1) confirms that KIF18A’s C-terminal tail is necessary to maintain microtubule association when the motor is inhibited by KBP, 2) suggests that the KIF18A motor domain is unable to associate with microtubules when bound to KBP, similar to findings for the kinesin-3 motor KIF1A (Kevenaar et al., 2016), and 3) supports the conclusion that KBP is cytosolic.

**Figure 5.**
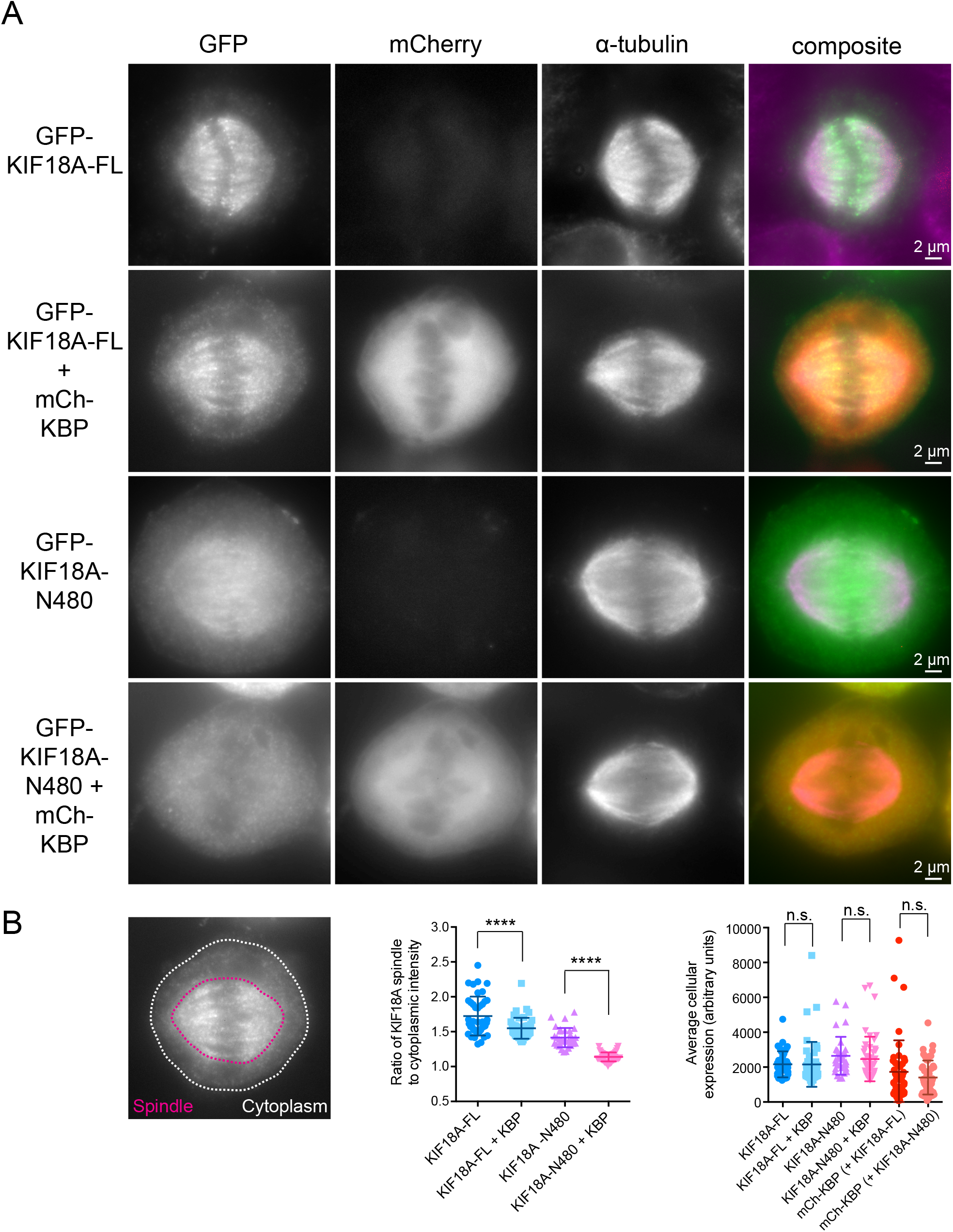
KIF18A’s C-terminal tail maintains spindle localization upon KBP overexpression. (**A**) Metaphase cells arrested in MG132 transfected with GFP-tagged KIF18A full-length (FL) or C-terminally truncated GFP-tagged KIF18A-N480 with or without mCh-KBP. (**B**) (*left*) Representative image indicating areas used to define spindle (red) and cytoplasmic (yellow) GFP-KIF18A fluorescence intensity. (*middle*) Quantification of background subtracted spindle-to-cytoplasmic ratio determined for individual cells in each condition. ****, adjusted p < 0.0001 with 95% confidence interval by one-way ANOVA with Tukey’s multiple comparisons test. (*right*) Quantification of background subtracted average fluorescence intensity of the indicated constructs n.s., not significant with 95% confidence interval by one-way ANOVA with Tukey’s multiple comparisons test. Data obtained from three independent data sets with the following cell numbers: GFP-KIF18A-FL (41), GFP-KIF18A-FL + mCh-KBP (47), GFP-KIF18A-N480 (32), GFP-KIF18A-N480 + mCh-KBP (47).

KIF15 contains two coiled-coil domains. The C-terminal coiled-coil is responsible for autoinhibition of the motor heads, whereas the proximal coiled-coil displays microtubule affinity and allows microtubule crosslinking and sliding in the spindle (Reinemann et al., 2017; Sturgill et al., 2014). Since KBP also alters KIF15 localization within the spindle, we compared the effects of KBP on the spindle localization of full-length (FL) KIF15 and a truncation mutant (KIF15-N700) containing only the proximal coiled-coil. Surprisingly, GFP-KIF15-FL co-expressed with mCherry-KBP localized stronger to the spindle than GFP-KIF15-FL alone (Figure 6B, left), and redistributed closer to the poles (Figure 6A), indicating that KBP may alter the equilibrium between the KIF15 autoinhibited and active binding states. GFP-Kif15-N700 displayed weak spindle localization in cells (Figure 6A) and became more cytosolic in the presence of mCherry-KBP (Figure 6B, middle). Taken together, these results suggest that KIF18A and KIF15 relocalize within the spindle via their C-termini when their motor domains are complexed with KBP.

**Figure 6.**
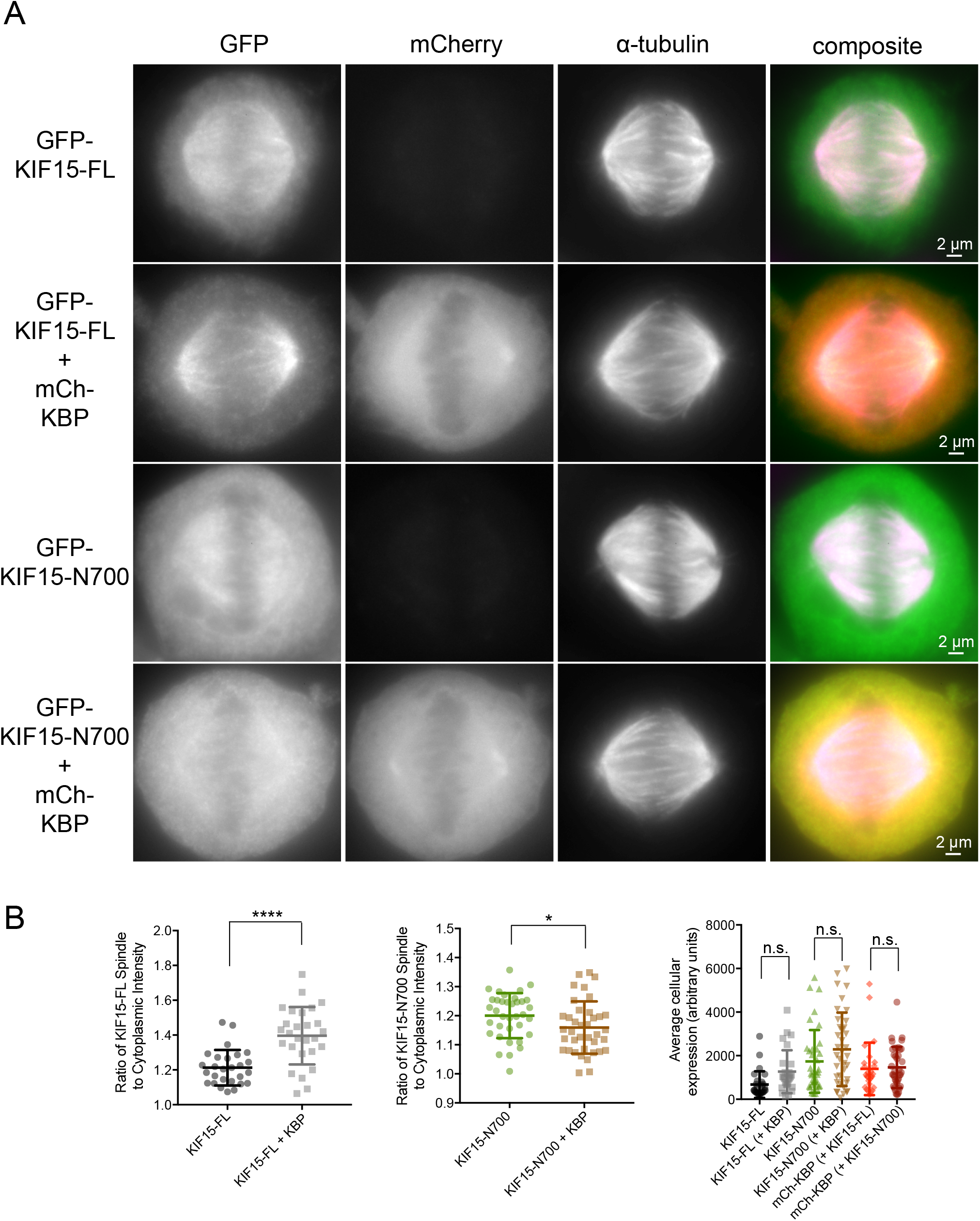
KIF15’s C-terminal tail contributes to spindle localization in the presence of KBP. (**A**) Metaphase cells arrested in MG132 transfected with GFP-tagged KIF15 full-length (FL) or C-terminally truncated GFP-tagged KIF15-N700 with or without mCh-KBP. (**B**) Quantification of background subtracted spindle-to-cytoplasmic ratio of GFP in individual cells expressing GFP-KIF15-FL (*left*) and GFP-KIF15-N700 (*middle)*. ****, p < 0.0001; *, p = 0.0387 by unpaired t-test. (*right*) Quantification of background subtracted average fluorescence intensity of the indicated constructs. n.s., not significant with 95% confidence interval by one-way ANOVA with Tukey’s multiple comparisons test. Data obtained from three independent data sets with the following cell numbers: GFP-KIF15-FL (27), GFP-KIF15-FL + mCh-KBP (27), GFP-KIF15-N700 (36), GFP-KIF15-N700 + mCh-KBP (38).

### KBP overexpression prevents KIF15-driven spindle stabilization

Given its ability to inhibit the motor activities and mitotic localizations of KIF18A and KIF15, we predicted that KBP would limit the functions of both kinesins in dividing cells. KIF15 has a partially overlapping role with Eg5 in maintaining spindle bipolarity (Tanenbaum et al., 2009), and is essential for bipolar spindle formation when Eg5 is inhibited (Sturgill et al., 2016; Tanenbaum et al., 2009). Thus, to determine the effects of KBP on KIF15’s spindle function, we treated cells expressing either mCherry alone or mCherry-KBP with the Eg5 inhibitor STLC or DMSO, as a negative control (Figure 7A). Fixed metaphase cells were scored for bipolar or monopolar spindles in each treatment condition (Figure 7B) and normalized to the DMSO control. Overexpression of mCherry-KBP induced a two-fold increase in the number of monopolar spindles, indicating that KBP indeed inhibits KIF15’s spindle stabilization function (Figure 7C). However, the effect of KBP overexpression was not as severe as complete inhibition of KIF15, which induces monopolar spindles in nearly 100% of mitotic cells in the presence of STLC (Sturgill et al., 2016).

**Figure 7.**
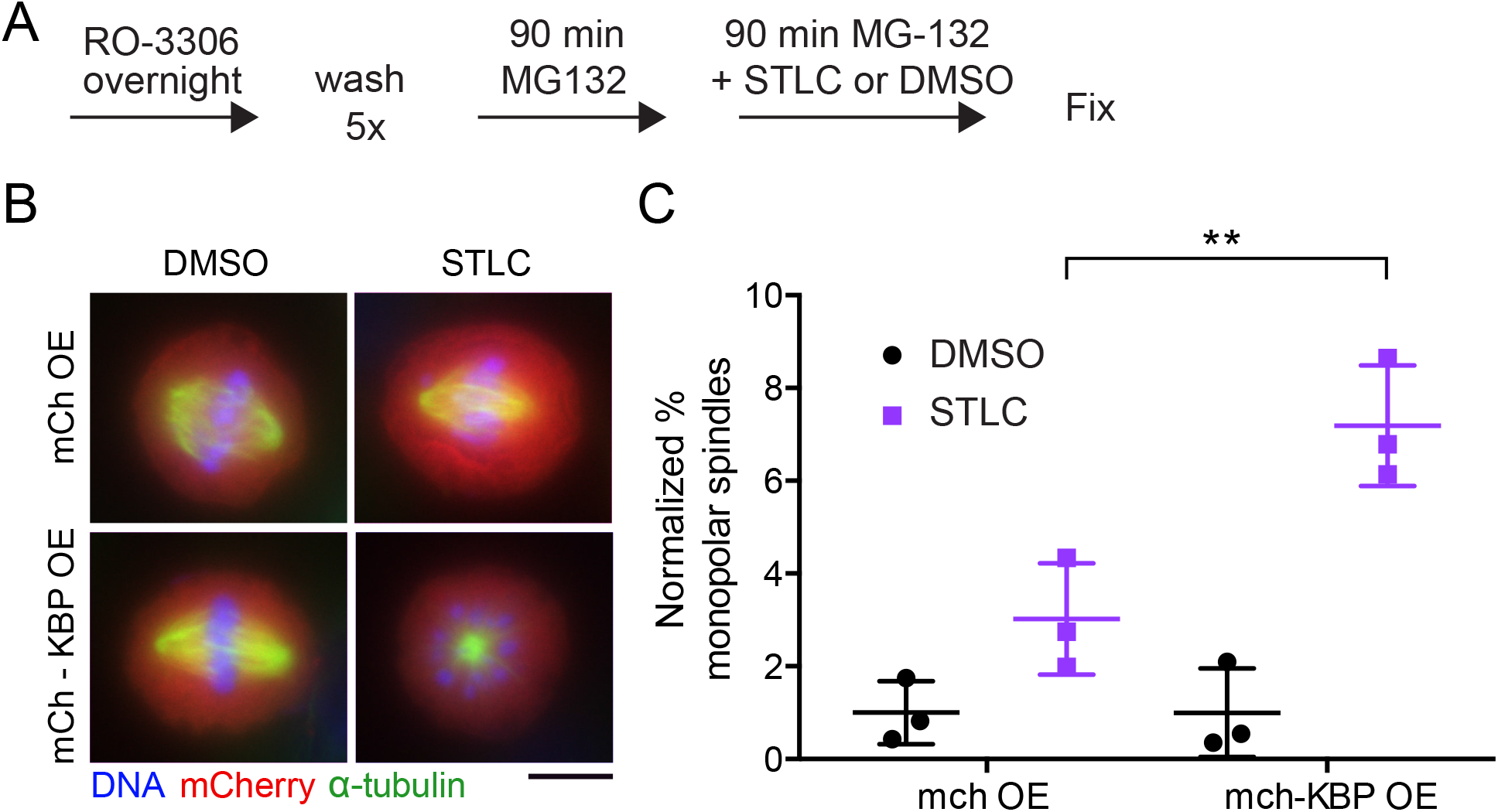
KBP overexpression inhibits KIF15-driven spindle stabilization. (**A**) Schematic of spindle collapse assay. (**B**) Metaphase cells expressing mCh only or mCh-KBP treated with DMSO or the Eg5 inhibitor STLC, scale bar is 10 μm. (**C**) Quantification of the percentage of mitotic cells with monopolar spindles, normalized to the DMSO condition. Data obtained from three independent experiments with the following cell numbers: mCh only DMSO (103), mCh only STLC (134), mCh-KBP DMSO (200), mCh-KBP STLC (269). **, adjusted p = 0.0058 with 95% confidence interval by two-way ANOVA with Tukey’s multiple comparisons test.

### Alteration in KBP expression influences kinetochore oscillations

KIF18A accumulates at kinetochore microtubule plus-ends and suppresses microtubule dynamics, which contributes to the confinement of chromosomes near the metaphase plate (Du et al., 2010; Stumpff et al., 2008). KIF18A knockdown increases the distance of kinetochores from the metaphase plate and the amplitude of kinetochore oscillations during metaphase, while KIF18A overexpression dampens oscillations (Stumpff et al., 2008). These effects can be assayed by measuring the deviation from the metaphase plate (DMP) and the deviation from average position (DAP) of kinetochores in live metaphase cells (Stumpff et al., 2008; Stumpff et al., 2012). To determine the effects of altered KBP expression on kinetochore oscillations, GFP-CENP-B expressing cells were transfected with control siRNAs, KBP siRNAs, or plasmids encoding mCherry or mCherry-KBP. Mitotic cells were then imaged using time-lapse microscopy (Videos S5-S8), and analyzed by tracking GFP-CENP-B labeled kinetochore pairs. Figures 8A and 8B show representative kinetochore pair tracks and kymographs for all conditions. KBP knockdown showed a significant reduction in both the DMP and DAP compared to control knockdown cells (Figure 8C, black data). In contrast, mCherry-KBP overexpression significantly increased both the DMP and DAP measurements compared to mCherry alone (Figure 8C, red data). These results are consistent with KBP inhibiting KIF18A’s function in controlling kinetochore oscillations and are in agreement with the measured effects of KBP on chromosome alignment (Figure 1, S2, and S3).

**Figure 8.**
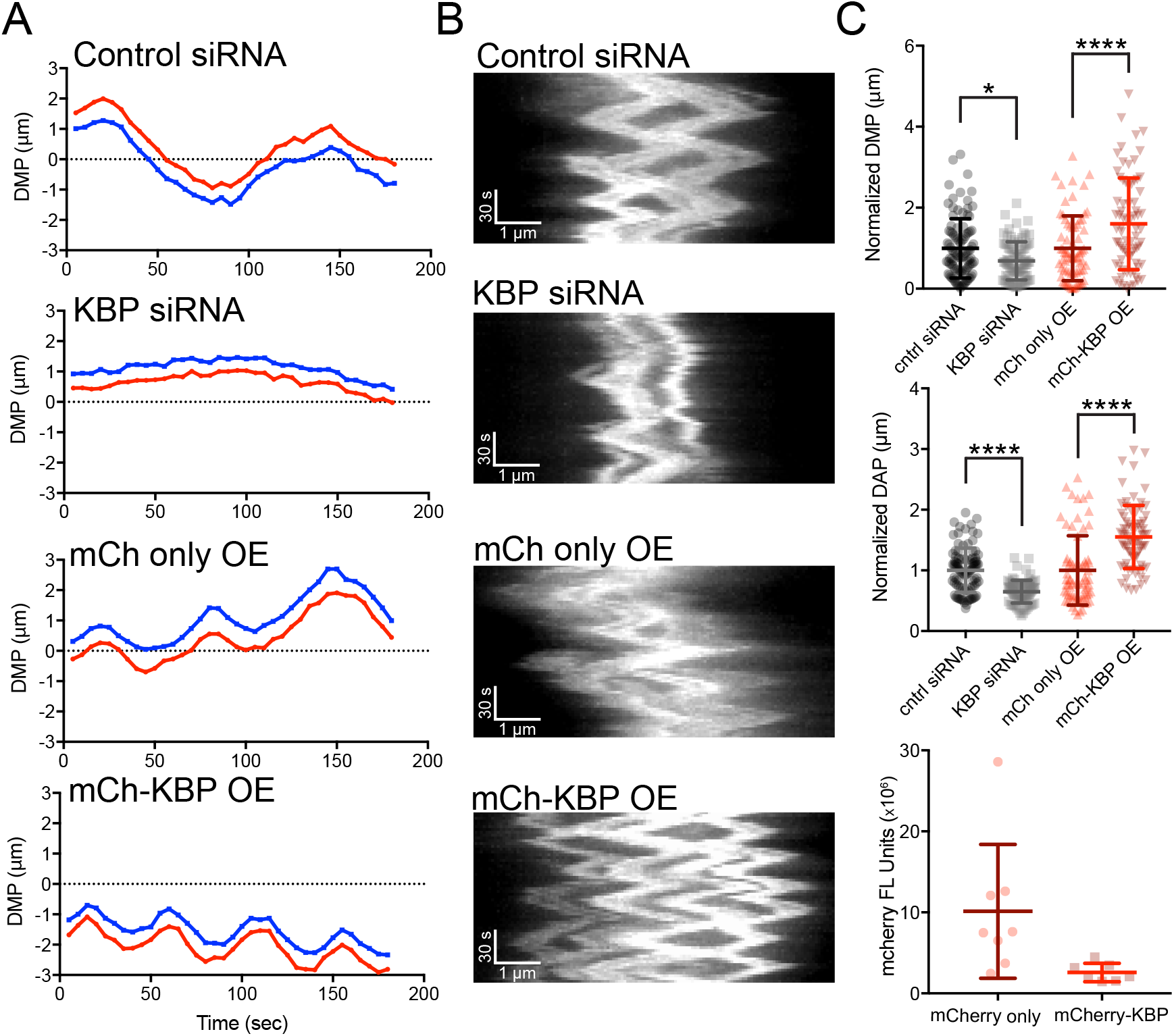
Altered KBP expression affects kinetochore oscillatory movements. (**A**) Example traces showing the movement over time of representative kinetochore pairs relative to the metaphase plate (dotted line) in cells from the indicated experimental conditions. (**B**) Representative kymographs of GFP-tagged CENP-B dynamics in metaphase cells transfected with the indicated siRNAs or plasmids. Full videos available in supplemental material (Videos S5-S8). (**C**) Quantification of the DMP (<u>d</u>eviation from the <u>m</u>etaphase <u>p</u>late), DAP (<u>d</u>eviation from <u>a</u>verage <u>p</u>osition), and mCh fluorescence. DMP and DAP values were normalized to control siRNA or mCh only conditions. *, adjusted p < 0.05; ****, adjusted p < 0.0001 with 95% confidence interval by one-way ANOVA with Tukey’s multiple comparisons test. Data obtained from the following numbers of cells over two or three independent experiments: control siRNA (10 (97 kinetochores)), KBP siRNA (12 (83 kinetochores)), mCh only (8 (69 kinetochores)), mCh-KBP (7 (75 kinetochores)).

### KBP knockdown or KIF18A/KIF15 overexpression results in lagging chromosomes during anaphase

Our data indicate that KBP limits the activity of KIF18A and KIF15 during mitosis, suggesting that KIF18A and KIF15 hyperactivity could be detrimental to spindle function and chromosome segregation. To test this, we analyzed chromosome segregation following KBP knockdown (Figure 9A). KBP depletion led to a two-fold increase in anaphase cells with lagging chromosomes compared to controls (Figure 9B). Importantly, of the cells that contained lagging chromosomes, those depleted of KBP exhibited a higher frequency of cells with more than one or two lagging chromosomes (Figure 9C). These results show that KBP is required for normal chromosome segregation.

**Figure 9.**
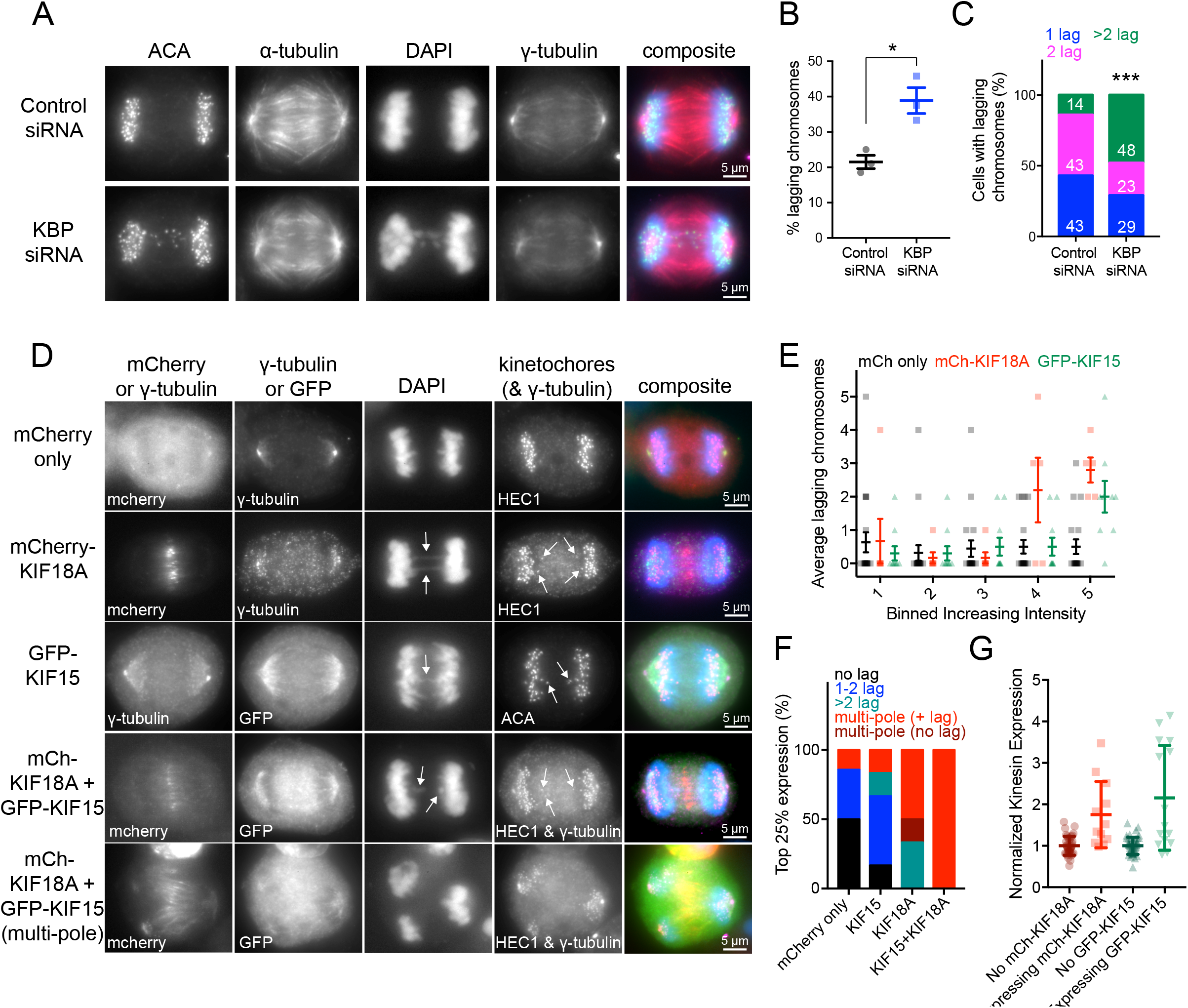
Cells deficient in KBP or overexpressing KIF18A and/or KIF15 display multiple lagging chromosomes in anaphase. (**A**) Anaphase cells treated with control or KBP siRNAs. ACA, anti-centromere antibodies. (**B**) Quantification of the percentage of lagging chromosomes under the two knockdown conditions. *, p = 0.0136 by unpaired t-test. Data obtained from three independent experiments with the following cell numbers: control siRNA (66), KBP siRNA (82). (**C**) Among the cells with lagging chromosomes, quantification of the number of lagging chromosomes per cell. 1 lagging chromosome, blue; 2 lagging chromosomes, pink; more than 2 lagging chromosomes, green. White numbers indicate percentage for each group. Number of cells with lagging chromosomes analyzed: control siRNA (13), KBP siRNA (31). ***, p < 0.001 by Chi-square analysis. (**D**) Anaphase cells overexpressing either mCherry only, mCherry-KIF18A, GFP-KIF15, or both mCherry-KIF18A and GFP-KIF15. Arrows point to lagging chromosomes. (**E**) Quantification of the average lagging chromosomes in cells as a function of mCh only, mCh-KIF18A, or GFP-KIF15 overexpression level. Background subtracted fluorescence intensity values were equally divided into bins. Individual data points are shown in background, foreground is average number of lagging chromosomes with standard deviations. Data obtained from seven independent experiments with the following cell numbers: mCh only (92), mCh-KIF18A (28), GFP-KIF15 (49). Significance: Bin 4 mCh only vs. mCh-KIF18A adjusted p = 0.0044; Bin 5 mCh only vs. mCh-KIF18A adjusted p < 0.0001; Bin 5 mCh only vs. GFP-KIF15 adjusted p = 0.0016 with 95% confidence interval by two-way ANOVA with Tukey’s multiple comparisons test. (**F**) Breakdown of the number of lagging chromosomes or multipolar spindle phenotypes observed in the 25% highest expressing cells for each condition. No lagging chromosomes, black; 1-2 lagging chromosomes, blue; more than 2 lagging chromosomes, green; multipolar spindles with lagging chromosomes, red without lagging chromosomes, dark red. Data obtained from four independent experiments with the following total cell numbers: mCh only (55), mCh-KIF18A (19), GFP-KIF15 (23), mCh-KIF18A and GFP-KIF15 (18). p < 0.0001 by Chi-square analysis. (**G**) Normalized kinesin expression levels in anaphase cells. Anaphase cells were imaged and partitioned into either expressing or not expressing mCh/GFP tagged kinesins. Plot displays background subtracted immunofluorescence levels from KIF18A or KIF15 specific antibodies normalized to the average from untransfected cells. Average fold increase in kinesin expression above endogenous for each kinesin: mCh-KIF18A (1.75), GFP-KIF15 (2.16). Data obtained from two independent data sets with the following cell numbers: KIF18A not expressing mCh-KIF18A (39), expressing mCh-KIF18A (12), KIF15 not expressing GFP-KIF15 (46), expressing GFP-KIF15 (15).

To determine if this phenotype is reflective of excess KIF18A and KIF15 motor activity, we overexpressed either mCherry alone, mCherry-tagged KIF18A, GFP-tagged KIF15, or mCherry-tagged KIF18A and GFP-tagged KIF15 in combination (Figure 9D). We observed that overexpression of either mCherry-KIF18A or GFP-KIF15 increased the frequency of lagging chromosomes and that the number of lagging chromosomes per cell scaled proportionally with increasing fluorescence intensity (Figure 9E). In addition, we found that cells with the highest expression (top 25%) of mCherry-KIF18A or GFP-KIF15 alone or in combination were much more likely to have multipolar spindles in addition to multiple lagging chromosomes, particularly in the double motor overexpression condition (Figure 9F). To determine the level of motor overexpression we labeled mCherry-KIF18A and GFP-KIF15 expressing cells with antibodies specific to the motors. We found that kinesin levels were increased ~1.5-fold on average compared to controls (Figure 9G), a range that is on par with measured protein variations that occur naturally within cell populations (Sigal et al., 2006). Taken together, these results indicate that KBP inhibition of KIF18A and KIF15 is necessary to prevent chromosome segregation errors during anaphase, which can result from relatively modest increases in kinesin levels.

## DISCUSSION

Our data indicate that KBP reduces the activities of KIF18A and KIF15 and that this control is required for proper mitotic spindle function and chromosome segregation. In agreement with published work (Kevenaar et al., 2016) we find that KBP blocks KIF18A and KIF15 activity by preventing their catalytic domains from interacting with the microtubule lattice. Accordingly, altered KBP expression in metaphase cells disrupts chromosome alignment, reduces spindle length, and destabilizes spindle bipolarity. These effects are consistent with inhibition of KIF18A and KIF15 function. The ability of KBP to disrupt motor-microtubule interactions is also supported by the effects of KBP overexpression on KIF18A and KIF15 localization. Excess KBP prevents KIF18A from accumulating at kinetochore microtubule plus-ends, instead causing the motor to indiscriminately bind spindle microtubules, most likely through a secondary microtubule-binding site in KIF18A’s C-terminal tail (Stumpff et al., 2011). When this region containing this site is removed, both KIF18A and KBP are exclusively cytosolic. Similarly, KBP inhibition of KIF15 causes the motor to adopt a more uniform distribution along the spindle, which depends on the KIF15 C-terminus (Sturgill et al., 2014). Phosphorylation has also been shown to alter KBP binding to kinesin motors (Kevenaar et al., 2016), and acetylation of KBP can promote its degradation (Donato et al., 2017), both of which could add to the complexity of KIF18A and KIF15 regulation. Our observation that full-length KIF15 localizes more strongly to the spindle when co-expressed with KBP, a finding we are not able to recapitulate *in vitro* with purified components, also implicates additional modes of regulation between KIF15 and KBP that should be explored in future studies.

Another recent report identified KBP as a regulator of KIF15 in mitotic cells. Our findings, however, contradict some of the primary conclusions of this study (Brouwers et al., 2017). Brouwers et al. reported that KBP promotes metaphase chromosome alignment by enhancing KIF15 localization near chromosomes and increasing kinetochore-microtubule stability (Brouwers et al., 2017). In contrast, we found that loss of KBP function leads to hyper-alignment of chromosomes, a phenotype consistent with increased KIF18A activity at kinetochore-microtubule plus-ends. Furthermore, our data indicate that KBP reduces the accumulation of KIF15 near the center of the spindle and that chromosomes align during metaphase despite complete loss of KIF15 activity, a conclusion supported by other studies of KIF15 function in mitosis (Sturgill and Ohi, 2013; van Heesbeen et al., 2014). Brouwers et al. also concluded that KBP localizes to the spindle. Evidence presented here and in previous studies (Kevenaar et al., 2016) suggests that KBP does not directly interact with microtubules. Our immunofluorescence analyses of endogenous KBP were complicated by non-specific spindle staining, but we did not observe spindle localization of mCherry-KBP. Thus, our data support the conclusion that KBP does not bind microtubules, but we cannot rule out that a small fraction of the protein localizes to spindles indirectly via kinesins that contain non-motor microtubule binding domains. Along these lines, the infrequent association of KBP with microtubules in interphase cells (Figure 1B) may be indirect, mediated through kinesin-2 and kinesin-3 motors.

Our finding that KBP has a greater inhibitory effect on KIF18A compared to KIF15 agrees with previous immunoprecipitation assays (Kevenaar et al., 2016). Differential preference for the two motors could explain the range of spindle lengths observed in Figure 1E, where KBP overexpression resulted in a cell population with longer spindle lengths on average even though knockdown of KIF18A and KIF15 in combination produced cells with normal spindle lengths (Figure 2C). The longer spindles in cells expressing low levels of mCherry-KBP could be caused by specific inhibition of KIF18A, while the shorter spindles in cells expressing higher levels of mCherry-KBP could result from KIF15 inhibition (Figure 1E, right). Interestingly, cells that expressed low levels of exogenous KBP but were not arrested in metaphase did not form long spindles (Figure S3B). This result is consistent with increased spindle length in KIF18A knockdown cells depending on mitotic arrest (Mayr et al., 2007). Why might KBP have evolved a higher potency for KIF18A? Our analyses of chromosome segregation suggest that cells are less tolerant of KIF18A overexpression. Therefore, increased specificity of KBP for KIF18A may be necessary to ensure accurate chromosome segregation.

The regulation of both KIF18A and KIF15 by KBP is required to prevent lagging chromosomes in anaphase. Given the multipolar phenotype observed from overexpression of KIF18A and KIF15, it is likely that these lagging chromosomes are caused, at least in part, by merotelic attachments. Merotelic attachments are a common cause of segregation errors, especially when the integrity of the spindle and control of chromosomal oscillations are compromised (Bakhoum and Compton, 2012; Cimini et al., 2001). Other mechanisms such as blocking error correction in kinetochore-microtubule attachment or linking neighboring kinetochore microtubules are also possible (Vladimirou et al., 2013). We have found that our kinesin overexpression assays at most increase the expression of KIF18A and KIF15 by four-fold compared to endogenous levels (Figure 9G). However, cells with the highest levels of kinesin expression exhibit multipolar spindles, which were not observed in KBP knockdown cells. Thus, we speculate that cells lacking KBP form bipolar spindles and lagging chromosomes due to increases in KIF18A and KIF15 activity that are lower than four-fold.

We propose a model where KBP functions as a protein buffer in the mitotic system, acting through its varying affinity for specific kinesin substrates as a referee to maintain an optimal level of KIF18A and KIF15 motor activity within the spindle. Why would a mode of regulation evolve to inhibit specific kinesin function when there are clearly established modes of transcriptional, translational, and ubiquitin-mediated degradation regulation (Nath et al., 2015)? Precedence for sequestration as a regulatory mechanism has been established for the actin and microtubule cytoskeletons, as well as enzymes such as cyclin-dependent kinases (Harper et al., 1993; Jourdain et al., 1997; Safer et al., 1991; Xiong et al., 1993). Similar to these systems, the mitotic spindle requires finely balanced control (Prosser and Pelletier, 2017), leaving little wiggle room for variations in activity levels. Protein concentration can vary within a cell population up to 30% and can change significantly over cell generations, as shown by tracking protein expression through multiple cell divisions (Sigal et al., 2006). While KBP itself would also be subject to this variability, there is already evidence that “excess” KBP, KBP unbound to kinesins, is targeted for degradation by acetylation (Donato et al., 2017). We propose that KBP functions to ensure that minor fluctuations in protein levels do not interfere with these precisely balanced forces, thus promoting robust and reproducible division of genetic material amongst protein expression variations within cells.

## MATERIALS AND METHODS

### Plasmids and siRNAs

mCherry-KBP and mCherry-KIF18A were constructed by Gateway Cloning Technology (Thermo Fisher Scientific). First, the KBP gene (HA-KBP plasmid received as a kind gift from Casper Hoogenraad (Kevenaar et al., 2016)) and KIF18A open reading frames were inserted into pCR8/GW/TOPO (Thermo Fisher Scientific). Correctly inserted clones were confirmed by sequencing. The KBP and KIF18A genes were then inserted by LR recombination reaction (Thermo Fisher Scientific) into a pCMV *ccdB* destination vector containing an N-terminal mCherry gene in frame with R1/R2 recombination sites. Clones were screened by restriction digestion, and the full open reading frames were confirmed by sequencing. For protein expression, GST-KBP was constructed by isothermal assembly, where the full-length human KBP cDNA (GE Healthcare, CloneId:4550085) was inserted into the pGEX6P1 vector (Amersham). Clones were screened by Sal1 and EcoR1 digestion and the full open reading frame confirmed by sequencing.

Other constructs used in this study are described elsewhere: GFP-CENP-B (Wordeman et al., 2007), GFP-KIF15 and GFP-KIF15-N700 (Sturgill et al., 2014), GFP-KIF18A-FL and GFP-KIF18A-N480 (Du et al., 2010; Stumpff et al., 2011), and CMV-mCherry alone previously used to clone mCherry-MCAK (Peris et al., 2009). Scrambled control siRNA was obtained from Thermo Fisher Scientific (Silencer Negative Control #2). KBP siRNA anti-sense sequence (Kevenaar et al., 2016): 5’-UAUCAUAGUAAGCAUGUGCUU-3’ (Qiagen). KIF18A siRNA anti-sense sequence (Kim et al., 2014; Stumpff et al., 2008; Stumpff et al., 2012): 5’-GCUGGAUUUCAUAAAGUGG-3’ (Ambion). KIF15 siRNA anti-sense sequence (Sturgill and Ohi, 2013; Tanenbaum et al., 2009): 5’-GGACAUAAAUUGCAAAUAC-3’ (Dharmacon). A SMARTpool of four siRNA sequences (Dharmacon) was used to target human KIF14.

### Cell Culture and Transfections

HeLa and RPE1 cells were cultured at 37°C with 5% CO_2_ in MEM-alpha medium (Life Technologies) containing 10% FBS plus 1% penicillin/streptomycin (Gibco). The HeLa KIF15 knockout cell line (Sturgill et al., 2016) was cultured in DMEM (Life Technologies) with the above supplements. For siRNA transfections in a 24-well format, approximately 80,000 cells in 500 μL MEM-alpha medium were treated with 35 pmol siRNA and 1.5 μL RNAiMax (Life Technologies), preincubated for 20 minutes in 62.5 μL OptiMeM (Life Technologies). Cells were treated with siRNAs for 48 hours before fixing or imaging. Plasmid transfections were conducted similarly, but with 375 ng kinesin or mCh-KBP plasmid DNA (100 ng GFP-CENP-B) and 2 μL LTX (Life Technologies). Plasmid transfections were incubated for 24 hours before fixing or imaging. Transfections were scaled up by well diameter as needed. For assays with sequential knockdown and plasmid transfections, the media was changed before plasmid transfection mix added.

### Western Blot Analysis

For KBP western blot detection, approximately 350,000 cells were plated into a 6-well dish and transfected as described above (scaled siRNA: 150 pmol siRNA + 6 μL RNAiMax in 250 uL OptiMeM; scaled plasmid: 1.5 μg plasmid + 8 μL LTX in 250 μL OptiMeM). Cells were washed once in phosphate buffered saline (PBS: Thermo Fisher Scientific) and lysed in 200 μL PHEM Lysis Buffer (60 mM PIPES, 10mM EGTA, 4mM MgCl_2_, 25mM HEPES pH 6.9) with 1% Triton-X and Halt protease and phosphatase inhibitors (Thermo Fisher Scientific) by scraping the plate and incubating on ice for 10 minutes. Then lysates were spun at maximum speed for 10 minutes and 180 μL supernatant added to 60 μL 4 X Laemmli Buffer prepared with βME (BioRad). Samples were run on precast 4-15% gradient gels (BioRad), transferred to low-fluorescence PVDF membrane (BioRad), blocked for 1 hour in 1:1 Odyssey blocking buffer (Li-Cor) and Tris Buffered Saline (TBS: 150 mM NaCl, and 50 mM Tris Base pH 7.4) + 0.1% TWEEN-20 (TBST). Primary antibodies against mouse-KBP (Abnova, 1:1,000) and rabbit-γ tubulin (Abcam, 1:1,000) were added in TBS overnight at 4°C. The membrane was washed with 2X TBST for 5 minutes each, then secondary antibodies (goat anti-mouse IgG DyLight 800 conjugate and donkey anti-rabbit IgG DyLight 680 conjugate, both at 1:10,000, Thermo Scientific) added in 1:1 Odyssey blocking buffer and TBST for 1 hour at room temperature. The membrane was washed with 2X TBST, then 1X TBS for 5 minutes each and developed on an Odyssey CLx (Li-Cor).

### Cell Fixation and Immunofluorescence

For metaphase observations, cells were treated with 20 μM MG132 (Selleck Chemicals) for 1-2 hours before fixation. MG132 treatment is indicated in figure legends. For spindle collapse assays, cells were collected in G2 overnight with 9 μM RO-3306 (Axxora), washed 5X with DMEM supplemented with FBS and pen/strep, then treated for 90 minutes with 5 μM MG132, followed by 90 minutes with 5 μM MG132 + 10 μM DMSO or 10 μM STLC (Sigma-Aldrich). Cells were fixed on coverslips in -20°C methanol (Fisher Scientific) plus 1% paraformaldehyde (Electron Microscopy Sciences) for 10 minutes on ice, dunked briefly and then washed 3X for 5 minutes each in TBS. Coverslips were blocked with 20% goat serum in antibody-diluting buffer (Abdil: TBS pH 7.4, 1% bovine serum albumin, 0.1% Triton-X, and 0.1% sodium azide) for 1 hr at room temperature before the primary antibodies diluted in Abdil were added for 1 hour at room temperature: mouse anti-α-tubulin at 1 μg/mL (Sigma-Aldrich), mouse anti-γ-tubulin at 1 μg/mL (Sigma-Aldrich), rabbit anti-KIF18A at 2 μg/mL (Bethyl), rabbit anti-KIF15 at 1 μg/mL (Sturgill and Ohi, 2013), rabbit anti-GFP at 4 μg/mL (Molecular Probes), rabbit anti-KIF14 at 2 μg/mL (Abcam), mouse anti-KBP at 1 μg/mL (Abnova). Cells were incubated with the following primary antibodies overnight at 4°C: human anti-centromere antibodies (ACA) at 2.5 μg/mL (Antibodies Incorporated), mouse anti-HEC1 at 0.5 μg/mL (GeneTex). All mCherry images are direct measurement of mCherry fluorescence. Incubation with 1 μg/mL goat secondary antibodies against mouse, rabbit, or human IgG conjugated to Alex Fluor 488, 594, or 647 (Molecular Probes) occurred for 1 hour at room temperature. Two 5-minute washes in TBS were performed between rounds of primary and secondary antibody incubations, finishing with three TBS washes before mounting with Prolong Gold anti-fade mounting medium with 4’,6-Diamidine-2’-phenylindole dihydrochloride (DAPI) (Molecular Probes).

### Protein Purification

His_6_-KIF15-N420, His_6_-KIF15-N700, His_6_-GFP-KIF15-FL, KIF18A-N480-GFP-His_6_, and KIF18A-FL-GFP-His_6_ purifications have been described previously (Reinemann et al., 2017; Stumpff et al., 2011; Sturgill et al., 2014).

GST-KBP was expressed in BL21DE3 cells with 0.4 mM IPTG overnight at 16°C. Cells were pelleted and flash frozen in liquid nitrogen and stored at -80°C. For purification, pellets were resuspended in lysis buffer (1xPBS, 500 mM NaCl, 5 mM β-mercaptoethanol (β-ME), 1% NP40 and protease inhibitors (Complete Protease Inhibitor Cocktail tablets, Sigma-Aldrich). Lysate was incubated with 1 mg/mL of lysozyme for 30 minutes on ice, followed by sonication. Lysate was clarified by centrifugation at 35,000 rpm for 1 hour at 4°C in a Ti 45 rotor (Beckman). Clarified supernatant was incubated with 2 mL of glutathione-agarose (Sigma-Aldrich) for 1 hour at 4°C and washed with ~200 mLs of wash buffer (1xPBS, 500 mM NaCl and 5 mM β-ME). Protein was eluted with 1M Tris (pH 8.0), 3M KCl and 200 mM glutathione and peak fractions were buffered exchanged into storage buffer (10 mM K-HEPES pH 7.7, 100 mM KCl and 1 mM DTT) using a PD 10 desalting column (GE Healthcare). Protein concentration was determined using Bradford assays and powdered sucrose was added 10% w/v before the protein was aliquoted, frozen in liquid nitrogen, and stored in -80°C.

### Microscopy

Cells were imaged on a Nikon Ti-E inverted microscope (Nikon Instruments) with Nikon objectives Plan Apo λ 60X 1.42NA and APO 100X 1.49 NA with a Spectra-X light engine (Lumencore) and environmental chamber. The following cameras were used: Clara cooled-CCD camera (Andor) and iXon X3 EMCCD camera (Andor). The Nikon Ti-E microscope is driven by NIS Elements software (Nikon Instruments). Gliding filament assays were performed on a Nikon Elements controlled Eclipse 90i (Nikon) equipped with a 60x 1.4 NA (Nikon) objective and a Cool Snap HQ2 CCD camera (Roper). Spindle collapse assay cells were acquired using a 60X 1.4 NA objective (Olympus) on a DeltaVision Elite imaging system (GE Healthcare) equipped with a Cool SnapHQ2 charge-coupled device (CCD) camera (Roper).

### Gliding Filament Assays

Flow chambers were assembled using a glass slide, a smaller No. 1.5 glass coverslip and double-sided tape to create a ~20 μl chamber. Approximately 1 μM of His_6_-KIF15-N700 or KIF18A-N480-GFP-His_6_ was incubated in the chamber for 3 minutes. The chamber was then washed with 60 μL of wash buffer (WB, [1x BRB80, 1 mM MgATP, 0.5 mg/mL casein]) and next incubated with 1% Pluronic F-127 for 1 minute to passivate the surface. The chamber was again washed with 60 uL of WB and then incubated with 0.5-1 μM of Alexa-594 labelled MTs (1:10, labeled:unlabeled tubulin) for 3 minutes. The chamber was then washed with 60 μL of flow cell buffer with or without GST-KBP (WB plus 70 mM β-ME, 0.035 mg/mL catalase, 0.2mg/mL glucose oxidase, 4.5 mg/mL glucose) and then imaged by epifluorescence microscopy. Time lapse sequences spanned 2 minutes with acquisitions captured every 2 seconds. ImageJ was used for image analysis and MT velocity was calculated by measuring the distance the MT travelled in total during the 2 minute movie (120 seconds). The average velocity of control slides from one experimental day was calculated and used to calculate the percent inhibition using the following equation,

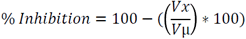

where *Vx* is the velocity of a single MT and *Vμ* is the average velocity of control MTs from that experimental day.

### Co-pelleting Assays

Microtubule co-pelleting was performed by incubating 700 nM of KIF18A-N480-GFP-His_6_ or 1 μM of His_6_-KIF15-N420 with 2 μM of GST-KBP in reaction buffer (10 mM K-HEPES pH 7.7, 50 mM KCl, 10 mM DTT, 1 mM MgCl_2_ and 20 μM taxol (Sigma-Aldrich)) for 10 minutes at room temperature to promote their interaction. KIF18A-N480-GFP-His_6_, His_6_-KIF15-N420, and GST-KBP were clarified by spinning at 90k rpm for 10 minutes at 4°C prior to use in assay. Next, 1 μM of taxol stabilized MTs and 1 mM AMPPNP (Sigma-Aldrich) was added to each reaction and incubated at room temperature for 15 minutes. The reaction was then spun over 150 μL of sucrose cushion (10 mM K-HEPES pH 7.7, 50 mM KCl, 20 μM taxol, 40% w/v sucrose) at 60k rpm for 20 minutes at 26°C. 50 μL of supernatant was removed and mixed 1:1 with 2x sample buffer (SB) (1x SB: 50 mM Tris-Cl, pH 6.8, 2% SDS, 6% glycerol, 1% βME and 200 μg/mL bromophenol blue). The remaining supernatant was removed and the supernatant/cushion interface was washed 2x with reaction buffer. The cushion was then removed and the pellet was gently washed with reaction buffer before being resuspended in 100 μl of 1x SB. Both supernatant and pellet samples from each reaction was boiled, vortexed and 30 μL was run on a 10% SDS-PAGE gel. The gel was then stained with Coomassie for ~30 minutes and destained overnight before imaging. ImageJ was used to quantify the integrated intensity of each protein band. Integrated intensity of the supernatant and pellet lanes for each reaction was combined to give yield a total protein integrated intensity value, which was used to calculate the percent of protein in the supernatant.

KIF18A-FL-GFP-His_6_ and His_6_-GFP-KIF15-FL co-pelleting experiments were performed as described above, except proteins were detected using Western blots instead of Coomassie. For KIF18A-FL-GFP-His_6_ experiments, 120 nM of kinesin and 342 nM of GST-KBP was used. For His_6_-GFP-KIF15-FL experiments, 135 nM of kinesin and 384 nM of GST-KBP was used. Blots were cut ~70 kDa and the top portion was probed with rabbit-His (MBL, 1:1000) and mouse-KBP (Abnova, 1:1000) and the bottom with α tubulin (DM1α, Abcam).

### Co-Immunoprecipitation Assays

Glutathione agarose slurry (Pierce) was washed with binding buffer (BB, 10 mM K-HEPES pH 7.7, 50 mM KCl, 1 mM DTT, 100 μM ATP, 0.1% NP-40) and incubated with 250 nM GST-KBP for 1 hour at 4°C with rotation. Agarose was then washed 3x with BB and incubated with 20 mg/mL BSA for 30 minutes at 4°C with rotation. Agarose was washed 3x with BB and incubated with 50 nM KIF18A-N480-GFP-His_6_, KIF18A-FL-GFP-His_6_, His_6_-KIF15-N420 or His_6_-GFP-KIF15-FL for 1 hour at 4°C with rotation.

Agarose was pelleted and supernatant was removed and saved, and the agarose pellet was resuspended in BB. Agarose was then washed 5-10x in BB. After washing, agarose was resuspended in 1x SB and boiled. Samples were loaded onto a TGX Stain-Free 10% Acrylamide gel (Bio-Rad), resolved by SDS-PAGE and transferred to a PVDF membrane for immunoblotting.

PVDF membranes were blocked with Odyssey blocking buffer (LI-COR Biosciences) diluted 1:1 in PBS for 1 hour at RT and probed with rabbit-His (MBL, 1:1000) and mouse-KBP (Abnova, 1:1000) overnight at 4°C. Membrane was washed 3x for 10 minutes with PBST and incubated with secondary antibodies conjugated to Alexa Fluor 700 (Invitrogen) or IRDye-800 (LI-COR) at 1:5000 for 1 hour at RT. Membrane was washed with PBST and PBS and bound antibodies were detected using an Odyssey fluorescence detection system (Mandel Scientific).

### Live Cell Imaging

MEM-alpha media was removed from transfected cells on poly-D-lysine coated filming dishes (MatTek) and replaced with CO_2_-independent imaging media (Gibco) supplemented with 10% FBS and 1% penicillin and streptomycin. Mitotic cells were identified by scanning for GFP-CENP-B fluorescence using the 60X objective and imaged with the 100X objective using 0.5 μm z-stacks every 5 seconds for approximately 5 minutes. After imaging, the XY position of the cell was marked and revisited within an hour of imaging to determine if the cell had divided for quality control. For mCherry or mCherry-KBP expressing cells, 0.5 μm z-stacks of the entire cell were taken in TRITC and GFP channels before time course imaging of GFP-CENP-B.

### Chromosome alignment and KIF15 localization analysis

Quantification of chromosome alignment was performed as described previously (Kim et al., 2014; Stumpff et al., 2012). Briefly, single focal plane images of metaphase cells were acquired with both spindle poles in focus. The Plot Profile command in ImageJ was used to measure the distribution of ACA-labeled kinetochore fluorescence within a region of interest (ROI) defined by the length of the spindle with a set height of 17.5 μm. The ACA signal intensity within the ROI was averaged along each pixel column, normalized, and plotted as a function of distance along the normalized spindle pole axis. These plots were analyzed by Gaussian fits using Igor Pro (Wavemetrics). The full width at half maximum intensity (FWHM) for the Gaussian fit and the spindle length are reported for each cell analyzed. KIF15 localization was performed similarly, but the endogenous KIF15 fluorescence was plotted across the spindle from a projection of 5 optical sections taken at 0.2 μm intervals and centered on the mid-cell plane. For chromosome hyperalignment quantification in RPE1 cells, the width of the metaphase plate was measured by determining the longitudinal distance between the most distal chromosomes as reported previously (Hafner et al., 2014). Lastly, for comparison with previously published effects of KBP knockdown on chromosome alignment (Brouwers et al., 2017), we analyzed full z-sections through non-arrested metaphase cells and scored for chromosomes off the metaphase plate.

### KIF18A line scan analysis

Fixed and stained cells were imaged at 100X with 0.2 μm z-stacks taken through the full cell. Line scans were manually measured using the α-tubulin and HEC1 fluorescent intensities to identify well-defined and locally isolated kinetochore microtubules. The profile intensities for α-tubulin, HEC1, and KIF18A were measured and recorded. Each channel was normalized internally to its highest value. Line scans were aligned in block by peak HEC1 intensity, and averaged for each pixel distance. Standard deviations are reported. Statistical comparisons were performed by fitting a linear regression of the accumulation slope (−0.8 μm to 0 μm) and comparing each condition to that of the control siRNA treatment using an F test.

### Chromosome oscillation analysis

Live kinetochore movements were tracked and quantified as reported previously (Stumpff et al., 2008; Stumpff et al., 2012). Briefly, z-slice projections of GFP-CENP-B movies were used to manually track individual kinetochore pairs with the MTrackJ plugin in ImageJ. Only kinetochore pairs near the middle of the cell were tracked; extreme peripheral kinetochore pairs were not included in the analysis. For mCherry expressing cells, a base cutoff of 1,000 background subtracted fluorescence units was required for inclusion in the data set. A macro was used to determine the position of the metaphase plate, which automatically fits a line to the thresholded GFP-CENP-B signal in a time-lapse projection. Kinetochore movement parameters were quantified from kinetochore tracks in Igor Pro (Wavemetrics) and reported as the deviation from the metaphase plate (DMP) and deviation from average position (DAP). Kymographs of GFP-CENP-B movements were created in ImageJ.

## SUPPLEMENTAL MATERIAL

Supplemental material includes: Figure S1 showing endogenous KBP antibody staining in HeLa and RPE1 cells; Figure S2 showing effects of KBP knockdown and overexpression in RPE1 cells; Figure S3 showing effects of KBP knockdown and overexpression in HeLa cells without MG132 treatment; Figure S4 showing that KIF14 knockdown does not affect chromosome alignment or spindle length and that KIF15 knockout cells have short spindles with aligned chromosomes; and Figure S5 showing velocity data from gliding assays in Figure 3 as well as co-IP and co-pelleting data from full-length motor constructs and an SDS-PAGE gel of purified proteins used in *in vitro* assays. Supplemental videos include example gliding filament movies for KIF15-N700 and KIF18A-N480 with and without KBP (Videos S1-4) and four examples of GFP-CENP-B kinetochore oscillations, one for each condition of control treatments, KBP siRNA, and mCh-KBP overexpression (Videos S5-8).

## ACKNOWLEDGMENTS

We thank Casper Hoogenraad’s group for the generous gift of their HA-KBP plasmid. We also thank Cindy Fonseca for technical support. This work was possible through the following funding sources: NIH R01 grants to J. Stumpff (R01GM121491) and R. Ohi (R01GM086610). Support also comes from a Susan G. Komen grant (CCR16377648) to J. Stumpff and R. Ohi was a Scholar of the Leukemia and Lymphoma Society. H. Malaby was supported by a Department of Defense PRCRP Horizon Award (W81XWH-17-1-0371). The authors declare no competing financial interests.

## AUTHOR CONTRIBUTIONS

H. Malaby, M. Dumas, R. Ohi, and J. Stumpff conceptualized the project and designed the experiments. H. Malaby, M. Dumas, and J. Stumpff performed the experiments. H. Malaby created the mCherry-KBP construct and performed the altered KBP expression experiments in metaphase and anaphase and kinesin overexpression in anaphase, and all KIF18A assays with the exception of the kinetochore oscillation experiments where she split the imaging with J. Stumpff. M. Dumas purified KBP and KIF15 proteins and performed the gliding filament, co-pelleting, and co-IP assays as well as KIF15 functional assays. H. Malaby and M. Dumas analyzed the data and assembled figures. H. Malaby and J. Stumpff wrote the manuscript. All authors reviewed and approved the final version of the manuscript.

## ABBREVIATIONS

KBP: Kinesin-Binding Protein
mCh: mCherry
OE: Overexpression
FWHM: Full width at half maximum

---

**Figure S1.**Analysis of KBP localization in HeLa and RPE1 cells. Metaphase HeLa (**A**) and RPE1 (**B**) cells from the indicated experimental conditions were arrested in MG132 and labeled with KBP antibodies. (**C**) (*left*) Quantification of mean, background subtracted KBP immunofluorescence in HeLa cells. Data normalized to mean for control siRNA condition. ****, p < 0.0001 by unpaired t-test. (*right*) Correlation plot of mCh-KBP fluorescence intensity vs. KBP immunofluorescence intensity in HeLa cells. Data obtained from one or two independent experiments with the following cell numbers: control siRNA (16), KBP siRNA (20), mCh-KBP OE (21). (**D**) (*left*) Quantification of mean, background subtracted KBP immunofluorescence intensity in RPE1 cells. Data normalized to mean for control siRNA condition. n.s., not significant by unpaired t-test. (*right*) Correlation plot of mCh-KBP fluorescence intensity vs. KBP immunofluorescence intensity in RPE1 cells. Data obtained from two independent experiments with the following cell numbers: control siRNA (45), KBP siRNA (61), mCh-KBP OE (25). (**E**) Western Blot against KBP showing protein levels under control or KBP siRNA treatment from RPE1 cells. γ-tubulin was used as a loading control.

**Figure S2.** KBP regulates chromosome alignment and spindle length in mitotic RPE1 cells. (**A**) Metaphase RPE1 cells arrested with MG132 following transfection with the indicated siRNAs or expression plasmids. ACA, anti-centromere antibodies. (**B**) Quantification of chromosome alignment (*left*) and spindle length (*right*) in cells treated with the indicated siRNAs. For RPE1 cell chromosome alignment, the metric of metaphase plate width (Hafner et al., 2014) was used instead of FWHM for increased sensitivity. ***, p = 0.0007; ****, p < 0.0001 by unpaired t-test. (*bottom*) Quantification of chromosome alignment (**C**) and spindle length (**D**) in mCh and mCh-KBP expressing cells. *, p = 0.0205; n.s., not significant by unpaired t-test. (*left*) Correlation plots of mCh only and mCh-KBP fluorescence intensity vs. FWHM or spindle length. Dotted lines are linear regression to show data trend. Data obtained from two (knockdown) or four (overexpression) independent experiments with the following cell numbers: control siRNA (38), KBP siRNA (55), mCh only (46), mCh-KBP OE (47).

**Figure S3.** KBP regulates mitotic chromosome alignment and spindle length in asynchronously dividing HeLa cells. (**A**) Metaphase HeLa cells transfected with control siRNA, KBP siRNA, or mCherry-KBP. No MG132 was added to media before fixation. ACA, anti-centromere antibodies. (**B**) Quantification of chromosome alignment (*top*) and spindle length (*bottom*) in metaphase cells from each experimental condition. The FWHM of kinetochore distribution in each cell was used as a metric for alignment as in Figure 1. Because chromosome compaction is less tight in unarrested metaphase cells, one standard deviation below the average FWHM in control siRNA cells was used as a cutoff for hyperalignment (3.7 um). ‡, p = 0.0266 by Chi-square analysis comparing hyperaligned populations. ****, adjusted p < 0.0001 with 95% confidence interval by one-way ANOVA analysis with Tukey’s multiple comparisons test of full data sets. Spindle length significance: **, adjusted p = 0.0011; ***, adjusted p = 0.0006 with 95% confidence interval by one-way ANOVA analysis with Tukey’s multiple comparisons test. (*right*) Correlation plot of mCh-KBP fluorescence intensity vs. FWHM or spindle length values. Dotted line is linear regression to show data trend. (**C**) An alternative assessment of chromosome alignment was carried out on same cells quantified in (B) by categorizing the percentage of cells containing one or more misaligned chromosomes. Complete optical sections were acquired for each cell to assess all chromosomes. n.s., not significant; ****, adjusted p < 0.0001 with 95% confidence interval by one-way ANOVA analysis with Tukey’s multiple comparisons test. Data obtained from three independent data sets with the following cell numbers: control siRNA (50), KBP siRNA (51), mCh-KBP OE (36).

**Figure S4.** KIF14 knockdown does not affect chromosome alignment or spindle length in metaphase and KIF15 knockout cells reproduce transient knockdown phenotypes. (**A**) Metaphase cells arrested in MG132 were labeled with KIF14 antibodies following treatment with control or KIF14 siRNAs. ACA, anti-centromere antibodies. (**B**) Quantification of chromosome alignment, spindle length, and (**C**) background subtracted KIF14 immunofluorescence intensity for both knockdown conditions. n.s., not significant; ****, p < 0.0001 by unpaired t-test. Data obtained from two independent data sets with the following cell numbers: control siRNA (45), KIF14 siRNA (60). (**D**) Untreated HeLa cells or KIF15 knockout (KO) HeLa cells arrested in metaphase were labeled with KIF15 antibodies. (**E**) Quantification of chromosome alignment and spindle length as in Figure 1. ***, p = 0.0002; ****, p < 0.0001 by unpaired t-test. Data obtained from two independent experiments with the following cell numbers: untreated HeLa (33), HeLa KIF15 KO (41).

**Figure S5.** Gliding filament velocities and characterization of KBP interaction with full-length kinesin motors. Microtubule gliding velocity distributions for (**A**) KIF18A-N480-GFP-His_6_ and (**B**) His_6_-KIF15-N700 in the presence of the indicated concentrations of GST-KBP. N values identical to Figure 3A. (**C**) Co-pelleting assay with KIF18A-FL-GFP-His_6_, GST-KBP, and/or microtubules (MTs). S, supernatant; P, pellet. Image is a Western blot probed with antibodies targeting 6xHis (α His), KBP (α KBP) and tubulin (DM1α). (**D**) Co-pelleting assay with KIF15-FL-GFP-His_6_, GST-KBP, and/or microtubules (MTs). S, supernatant; P, pellet. Image is a Western blot probed with antibodies targeting 6xHis (α His), KBP (α KBP) and tubulin (DM1α). (**E**) Co-immunoprecipitation (co-IP) of purified KIF18A-FL-GFP-His_6_ with GST-KBP. Glutathione agarose was first incubated with 250 nM GST-KBP and then with 50nM of KIF18A-FL-GFP-His_6_. S, supernatant; P, pellet. Image is a Western blot probed with antibodies targeting 6xHis (α His) and KBP (α KBP). (**F**) Co-immunoprecipitation (co-IP) of purified His_6_-GFP-KIF15-FL with GST-KBP. Glutathione agarose was incubated with 250 nM GST-KBP and then incubated with 50 nM of His_6_-GFP-KIF15-FL. S, supernatant; P, pellet. Image is a Western blot probed with antibodies targeting 6xHis (α His) and KBP (α KBP). (**G**) Purification of kinesin and KBP proteins. Coomassie stained gel of proteins purified for this study: His_6_-KIF15-N420 (MW = 45 kDa), His_6_-KIF15-N700 (MW = 76 kDa), KIF18A-N480-GFP-His_6_ (MW = 82 kDa) and GST-KBP (MW = 97 kDa). Molecular weight markers are indicated in kDa.

**Video S1. KIF18A gliding filament assay from Figure 3.** Time-lapse imaging of Alexa-594 labelled microtubules gliding along surface-immobilized KIF18A-N480-GFP-His_6_ (1 μM). Timestamps and scale bar included in the video.

**Video S2. KIF18A gliding filament assay with KBP from Figure 3.** Time-lapse imaging of Alexa-594 labelled microtubules gliding along surface-immobilized KIF18A-N480-GFP-His_6_ (1 μM) in the presence of 250 nM GST-KBP. Timestamps and scale bar included in the video.

**Video S3. KIF15 gliding filament assay from Figure 3.** Time-lapse imaging of Alexa-594 labelled microtubules gliding along surface-immobilized His_6_-KIF15-N700 (1 μM). Timestamps and scale bar included in the video.

**Video S4. KIF15 gliding filament assay with KBP from Figure 3.** Time-lapse imaging of Alexa-594 labelled microtubules gliding along surface-immobilized His_6_-KIF15-N700 (1 μM) in the presence of 250 nM GST-KBP. Timestamps and scale bar included in the video.

**Video S5. Kinetochore oscillations in control siRNA treated cells from Figure 6.** Time-lapse imaging of GFP-CENP-B expressing HeLa cell treated with control siRNAs. Timestamps and scale bar included in the video.

**Video S6. Kinetochore oscillations in KBP siRNA treated cells from Figure 6.** Time-lapse imaging of GFP-CENP-B expressing HeLa cell treated with KBP siRNAs. Timestamps and scale bar included in the video.

**Video S7. Kinetochore oscillations in mCh only expressing cells from Figure 6.** Time-lapse imaging of GFP-CENP-B expressing HeLa cell transfected with mCherry control plasmid. Timestamps and scale bar included in the video.

**Video S8. Kinetochore oscillations in mCh-KBP expressing cells from Figure 6.** Time-lapse imaging of GFP-CENP-B expressing HeLa cell transfected with mCherry-KBP. Timestamps and scale bar included in the video.

## REFERENCES

Alves, M.M., G. Burzynski, J.M. Delalande, J. Osinga, A. van der Goot, A.M. Dolga, E. de Graaff, A.S. Brooks, M. Metzger, U.L. Eisel, I. Shepherd, B.J. Eggen, and R.M. Hofstra. 2010. KBP interacts with SCG10, linking Goldberg-Shprintzen syndrome to microtubule dynamics and neuronal differentiation. Hum Mol Genet. 19:3642–3651.

Bakhoum, S.F., and D.A. Compton. 2012. Kinetochores and disease: keeping microtubule dynamics in check! Current opinion in cell biology. 24:64–70.

Brooks, A.S., A.M. Bertoli-Avella, G.M. Burzynski, G.J. Breedveld, J. Osinga, L.G. Boven, J.A. Hurst, G.M. Mancini, M.H. Lequin, R.F. de Coo, I. Matera, E. de Graaff, C. Meijers, P.J. Willems, D. Tibboel, B.A. Oostra, and R.M. Hofstra. 2005. Homozygous nonsense mutations in KIAA1279 are associated with malformations of the central and enteric nervous systems. Am J Hum Genet. 77:120–126.

Brouwers, N., N. Mallol Martinez, and I. Vernos. 2017. Role of Kif15 and its novel mitotic partner KBP in K-fiber dynamics and chromosome alignment. PloS one. 12:e0174819.

Cimini, D., B. Howell, P. Maddox, A. Khodjakov, F. Degrassi, and E.D. Salmon. 2001. Merotelic kinetochore orientation is a major mechanism of aneuploidy in mitotic mammalian tissue cells. The Journal of cell biology. 153:517–527.

Donato, V., M. Bonora, D. Simoneschi, D. Sartini, Y. Kudo, A. Saraf, L. Florens, M.P. Washburn, M. Stadtfeld, P. Pinton, and M. Pagano. 2017. The TDH-GCN5L1-Fbxo15-KBP axis limits mitochondrial biogenesis in mouse embryonic stem cells. Nat Cell Biol. 19:341–351.

Drevillon, L., A. Megarbane, B. Demeer, C. Matar, P. Benit, A. Briand-Suleau, V. Bodereau, J. Ghoumid, M. Nasser, X. Decrouy, M. Doco-Fenzy, P. Rustin, D. Gaillard, M. Goossens, and I. Giurgea. 2013. KBP-cytoskeleton interactions underlie developmental anomalies in Goldberg-Shprintzen syndrome. Hum Mol Genet. 22:2387–2399.

Du, Y., C.A. English, and R. Ohi. 2010. The kinesin-8 Kif18A dampens microtubule plus-end dynamics. Current biology: CB. 20:374–380.

Hafner, J., M.I. Mayr, M.M. Mockel, and T.U. Mayer. 2014. Pre-anaphase chromosome oscillations are regulated by the antagonistic activities of Cdk1 and PP1 on Kif18A. Nature communications. 5:4397.

Harper, J.W., G.R. Adami, N. Wei, K. Keyomarsi, and S.J. Elledge. 1993. The p21 Cdk-interacting protein Cip1 is a potent inhibitor of G1 cyclin-dependent kinases. Cell. 75:805–816.

Jourdain, L., P. Curmi, A. Sobel, D. Pantaloni, and M.F. Carlier. 1997. Stathmin: a tubulin-sequestering protein which forms a ternary T2S complex with two tubulin molecules. Biochemistry. 36:10817–10821.

Kevenaar, J.T., S. Bianchi, M. van Spronsen, N. Olieric, J. Lipka, C.P. Frias, M. Mikhaylova, M. Harterink, N. Keijzer, P.S. Wulf, M. Hilbert, L.C. Kapitein, E. de Graaff, A. Ahkmanova, M.O. Steinmetz, and C.C. Hoogenraad. 2016. Kinesin-Binding Protein Controls Microtubule Dynamics and Cargo Trafficking by Regulating Kinesin Motor Activity. Current biology: CB. 26:849–861.

Kim, H., C. Fonseca, and J. Stumpff. 2014. A unique kinesin-8 surface loop provides specificity for chromosome alignment. Molecular biology of the cell. 25:3319–3329.

Maliga, Z., M. Junqueira, Y. Toyoda, A. Ettinger, F. Mora-Bermudez, R.W. Klemm, A. Vasilj, E. Guhr, I. Ibarlucea-Benitez, I. Poser, E. Bonifacio, W.B. Huttner, A. Shevchenko, and A.A. Hyman. 2013. A genomic toolkit to investigate kinesin and myosin motor function in cells. Nat Cell Biol. 15:325–334.

Mayr, M.I., S. Hummer, J. Bormann, T. Gruner, S. Adio, G. Woehlke, and T.U. Mayer. 2007. The human kinesin Kif18A is a motile microtubule depolymerase essential for chromosome congression. Current biology: CB. 17:488–498.

Mayr, M.I., M. Storch, J. Howard, and T.U. Mayer. 2011. A non-motor microtubule binding site is essential for the high processivity and mitotic function of kinesin-8 Kif18A. PloS one. 6:e27471.

Nath, S., D. Ghatak, P. Das, and S. Roychoudhury. 2015. Transcriptional control of mitosis: deregulation and cancer. Front Endocrinol (Lausanne). 6:60.

Peris, L., M. Wagenbach, L. Lafanechere, J. Brocard, A.T. Moore, F. Kozielski, D. Job, L. Wordeman, and A. Andrieux. 2009. Motor-dependent microtubule disassembly driven by tubulin tyrosination. The Journal of cell biology. 185:1159–1166.

Prosser, S.L., and L. Pelletier. 2017. Mitotic spindle assembly in animal cells: a fine balancing act. Nature reviews. Molecular cell biology. 18:187–201.

Reinemann, D.N., E.G. Sturgill, D.K. Das, M.S. Degen, Z. Voros, W. Hwang, R. Ohi, and M.J. Lang. 2017. Collective Force Regulation in Anti-parallel Microtubule Gliding by Dimeric Kif15 Kinesin Motors. Current biology: CB. 27:2810–2820 e2816.

Safer, D., M. Elzinga, and V.T. Nachmias. 1991. Thymosin beta 4 and Fx, an actin-sequestering peptide, are indistinguishable. The Journal of biological chemistry. 266:4029–4032.

Sigal, A., R. Milo, A. Cohen, N. Geva-Zatorsky, Y. Klein, Y. Liron, N. Rosenfeld, T. Danon, N. Perzov, and U. Alon. 2006. Variability and memory of protein levels in human cells. Nature. 444:643–646.

Stumpff, J., Y. Du, C.A. English, Z. Maliga, M. Wagenbach, C.L. Asbury, L. Wordeman, and R. Ohi. 2011. A tethering mechanism controls the processivity and kinetochore-microtubule plus-end enrichment of the kinesin-8 Kif18A. Molecular cell. 43:764–775.

Stumpff, J., G. von Dassow, M. Wagenbach, C. Asbury, and L. Wordeman. 2008. The kinesin-8 motor Kif18A suppresses kinetochore movements to control mitotic chromosome alignment. Developmental cell. 14:252–262.

Stumpff, J., M. Wagenbach, A. Franck, C.L. Asbury, and L. Wordeman. 2012. Kif18A and chromokinesins confine centromere movements via microtubule growth suppression and spatial control of kinetochore tension. Developmental cell. 22:1017–1029.

Sturgill, E.G., D.K. Das, Y. Takizawa, Y. Shin, S.E. Collier, M.D. Ohi, W. Hwang, M.J. Lang, and R. Ohi. 2014. Kinesin-12 Kif15 targets kinetochore fibers through an intrinsic two-step mechanism. Current biology: CB. 24:2307–2313.

Sturgill, E.G., S.R. Norris, Y. Guo, and R. Ohi. 2016. Kinesin-5 inhibitor resistance is driven by kinesin-12. The Journal of cell biology. 213:213–227.

Sturgill, E.G., and R. Ohi. 2013. Kinesin-12 differentially affects spindle assembly depending on its microtubule substrate. Current biology: CB. 23:1280–1290.

Tanenbaum, M.E., L. Macurek, A. Janssen, E.F. Geers, M. Alvarez-Fernandez, and R.H. Medema. 2009. Kif15 cooperates with eg5 to promote bipolar spindle assembly. Current biology: CB. 19:1703–1711.

van Heesbeen, R.G., M.E. Tanenbaum, and R.H. Medema. 2014. Balanced activity of three mitotic motors is required for bipolar spindle assembly and chromosome segregation. Cell Rep. 8:948–956.

Vladimirou, E., N. McHedlishvili, I. Gasic, J.W. Armond, C.P. Samora, P. Meraldi, and A.D. McAinsh. 2013. Nonautonomous movement of chromosomes in mitosis. Developmental cell. 27:60–71.

Weaver, L.N., S.C. Ems-McClung, J.R. Stout, C. LeBlanc, S.L. Shaw, M.K. Gardner, and C.E. Walczak. 2011. Kif18A uses a microtubule binding site in the tail for plus-end localization and spindle length regulation. Current biology: CB. 21:1500–1506.

Wordeman, L., M. Wagenbach, and G. von Dassow. 2007. MCAK facilitates chromosome movement by promoting kinetochore microtubule turnover. The Journal of cell biology. 179:869–879.

Wozniak, M.J., M. Melzer, C. Dorner, H.U. Haring, and R. Lammers. 2005. The novel protein KBP regulates mitochondria localization by interaction with a kinesin-like protein. BMC Cell Biol. 6:35.

Xiong, Y., G.J. Hannon, H. Zhang, D. Casso, R. Kobayashi, and D. Beach. 1993. p21 is a universal inhibitor of cyclin kinases. Nature. 366:701–704.

